# A comprehensive genealogy of the replication associated protein of CRESS DNA viruses reveals a single origin of intron-containing Rep

**DOI:** 10.1101/687855

**Authors:** Lele Zhao, Erik Lavington, Siobain Duffy

**Affiliations:** Department of Ecology, Evolution and Natural Resources, School of Environmental and Biological Sciences, Rutgers, the State University of New Jersey

## Abstract

Abundant novel circular Rep-encoding ssDNA viruses (CRESS DNA viruses) have been discovered in the past decade, prompting a new appreciation for the ubiquity and genomic diversity of this group of viruses. Although highly divergent in the hosts they infect or are associated with, CRESS DNA viruses are united by the homologous replication-associated protein (Rep). An accurate genealogy of Rep can therefore provide insights into how these diverse families are related to each other. We used a dataset of eukaryote-associated CRESS DNA RefSeq genomes (n=926), which included representatives from all six established families and unclassified species. To assure an optimal Rep genealogy, we derived and tested a bespoke amino acid substitution model (named CRESS), which outperformed existing protein matrices in describing the evolution of Rep. The CRESS model-estimated Rep genealogy resolved the monophyly of *Bacilladnaviridae* and the reciprocal monophyly of *Nanoviridae* and the alpha-satellites when trees estimated with general matrices like LG did not. The most intriguing, previously unobserved result is a likely single origin of intron-containing Reps, which causes several geminivirus genera to group with *Genomoviridae* (bootstrap support 55%, aLRT SH-like support 0.997, 0.91-0.997 in trees estimated with established matrices). This grouping, which eliminates the monophyly of *Geminiviridae,* is supported by both domains of Rep, and appears to be related to our use of all RefSeq Reps instead of subsampling to get a smaller dataset. In addition to producing a trustworthy Rep genealogy, the derived CRESS matrix is proving useful for other analyses; it best fit alignments of capsid protein sequences from several CRESS DNA families and parvovirus NS1/Rep sequences.

## Introduction

Our understanding of eukaryotic circular Rep-encoding single-stranded DNA (CRESS DNA) viruses is changing, as this group is no longer restricted to plant and livestock infecting pathogens, but is now considered ubiquitous. With the application of the highly processive phi29 polymerase to enrich for circular DNA through rolling circle amplification, numerous publications have found CRESS DNA viruses, including on all continents (Haible et al., 2006; Inoue-Nagata et al., 2004; Li et al., 2010; Rosario et al., 2009, 2012; Wyant et al., 2012). Many CRESS DNA viruses have been associated with animals such as insects, birds, rodents, bats, chimps and humans, and in myriad environmental and animal tissue samples (reviewed in Zhao et al., 2019). Alongside the discovery driven, accelerated accumulation of new viral sequences deposited to GenBank, we are seeing commensurate taxonomical revisions and proposals to change the groupings of CRESS DNA viruses (Kazlauskas et al., 2017; Krupovic et al., 2016; Rosario et al., 2017; Varsani and Krupovic, 2017; Varsani and Krupovic, 2018; Varsani et al., 2014a; Varsani et al., 2014b; Varsani et al., 2017). The proposal of several new CRESS DNA viral families is anticipated (Kazlauskas et al., 2018)Abbas et al 2019) and additional unclassified sequences await the discovery of similar sequences before classification may be attempted. While we understand more about the diversity within this group than ever before, the relationships among CRESS DNA viral species, and even their classified families, are not well understood.

Building a phylogenetic tree is an established way to put new viral sequences into the context of well-characterized viruses. However, unlike eukaryotic cytochrome c oxidase I and prokaryotic 16S phylogenetic trees, viruses do not share a universal gene that can be used to reconstruct their evolutionary history. Instead, the deep phylogeny of viruses is typically restricted to groups that share at least one homologous protein. RNA dependent RNA polymerase is the shared gene used to study the relationships among RNA viruses (Koonin, 1991; Koonin and Dolja, 2012; Payne, 2017). Glycoprotein B and DNA polymerase protein sequences have been used for phylogenetic analyses and taxonomic classification of dsDNA herpesviruses (Chmielewicz et al., 2003; Ehlers et al., 1999; McGeoch and Gatherer, 2005). Despite varying genomic structure and size, all CRESS DNA viruses, by definition, share a replication associated protein (Rep) sequence, facilitating its use for development of a phylogeny of CRESS DNA viruses. While Rep gene sequences can be useful for studying the relationships within an individual family of CRESS DNA viruses (Simmonds et al., 2017), ssDNA viruses are known to evolve as quickly as RNA viruses (Duffy and Holmes, 2008; Duffy et al., 2008; Firth et al., 2009; Harkins et al., 2009; Shackelton et al., 2005), quickly saturating the information in their nucleotide sequences (Melcher 2010). Therefore, it is necessary to use protein sequences to determine the evolutionary relationships of divergent families and representatives of CRESS DNA viruses.

Unfortunately, methods available for describing CRESS viral protein evolution are not ideal. Previous attempts at building Rep trees have used a variety of amino acid substitution matrices, all of which may be poorly parameterized for CRESS DNA viruses. General matrices (such as LG, WAG, BLOSUM62, VT and PAM) are either estimated from aged and short protein sequence alignments, or from datasets containing protein sequences of multiple biological sources (mostly cellular). Organismal biologists in several areas have noticed the non-specific performance of these general matrices, and compensated for this inadequacy by estimating substitution matrices from highly specific protein sequences and then constructing phylogenies. Thus, the amino acid substitution matrices rtREV, cpREV, mtREV, HIVb, HIVw and FLU were developed from retro-transcribing viruses, chloroplasts, mitochondria, HIV and influenza sequences respectively (Adachi and Hasegawa, 1996; Adachi et al., 2000; Dang et al., 2010; Dimmic et al., 2002; Nickle et al., 2007b). Unsurprisingly, these specific matrices repeatedly outperform the general matrices in describing their designated sequences. The rtREV matrix has even been selected as the best-fitting matrix by ProtTest for CRESS DNA viral sequences on occasion (Dayaram et al., 2016; Dayaram et al., 2015a; Dayaram et al., 2015b; Kraberger et al., 2015b; Rosario et al., 2015). This result highlights the shortfalls of the general substitution matrices to describe CRESS DNA viral evolution rather than suggesting that CRESS DNA viruses evolve in an identical manner to retro-transcribing viruses.

To best understand the evolution of CRESS DNA viruses, we estimated and validated an amino acid matrix specific to eukaryotic-associated CRESS DNA viruses (which have a more recent common ancestor than they do with prokaryotic ssDNA viruses, Koonin and Ilyina, 1993). Sequences of the homologous protein Rep were used since it is necessarily present in all CRESS DNA viruses. As expected, this CRESS matrix outperformed all other established matrices in describing CRESS Rep phylogeny, and its Rep genealogy had several differences compared to trees estimated from the same alignment but using established protein matrices. However, all of our trees supported a previously unobserved relationship: a common origin for the intron-containing form of the Rep among some members of *Geminiviridae* and *Genomoviridae*. This clade destroys the reciprocal monophyly between these related families; our Rep genealogy supports monophyly for only three of the six established families of CRESS DNA viruses.

## Results

### Generation of a CRESS DNA virus Rep-specific amino acid substitution matrix

926 Rep sequences from eukaryotic-infecting or eukaryotic-associated CRESS DNA viruses were aligned, trimmed, and an initial maximum likelihood tree was built with the best-fitting model chosen in ProtTest3 (VT+G+F, uniformly chosen by AIC, AICc and BIC scores). The trimmed alignment was jackknifed ten times and the tree split accordingly. Each training set half was used for amino acid substitution matrix estimation in four ways: with HyPhy or FastMG, and starting with the seed matrix of VT or LG (a total of 40 estimated matrices). The four estimated matrices for each training dataset along with VT, LG and rtREV were fitted to the appropriate test set half and tree in PAML and goodness of fit was compared through maximum likelihood scores (Supplementary File 1). The FastMG estimated matrices always outperformed the established matrices, but the HyPhy estimated matrices did not.

We then evaluated the consistency of jackknifed matrices by comparing their Pearson correlation coefficients (Supplementary File 2). Matrices estimated by FastMG are very similar to each other, regardless of starting from the VT or LG matrix, with correlation coefficients ranging from 0.989 to 0.999. The HyPhy fit algorithm produced more varied matrices with correlation coefficients ranging from 0.824 to 1. The lowest correlations (as low as 0.79) were seen when comparing the three established matrices (VT, LG and rtREV) to the estimated matrices, substantiating that these matrices may not model the evolution of the Rep protein very well. Control analyses showed that the size of our training datasets was sufficient to reconstruct an established amino acid substitution matrix (Pearson correlation coefficients of the fmg matrices’ rates to WAG rates ranged from 0.93 to 0.97) without systematic over- or under-representation of substitution rates, as shown by the lack of correlation among matrices derived at different simulated levels of substitution rate variation (Supplementary File 3). This validates the potential for our 40 estimated matrices to accurately capture the patterns of protein evolution in CRESS DNA viral Reps.

### Matrix selection and comparison to established matrices

We selected one single trained matrix from the ten that best fit each of the test datasets to be the single matrix for all downstream analyses. We built maximum likelihood trees from ten test set alignments using the best performing matrix of every jackknifed training sets. Then, we rank ordered these ten matrices by the likelihood scores of the trees they generated (Supplementary File 4). The best matrix with the lowest sum in ranking (fmgVT-10) was named the CRESS matrix and used to construct the CRESS DNA viral Rep tree.

We compared the substitution rates of the CRESS matrix to rtREV, LG and VT, shown as heat maps are the log_10_ ratio of amino acid substitution (Figure 1). The CRESS matrix has generally lower rates of substitution compared to rtREV, has similar rates compared to LG and higher rates compared to VT. The CRESS matrix seems to have a consistently low proline (P) – Isoleucine (I) interchange rate compared to all other three matrices. Compared to VT, the CRESS matrix has higher rates of substitution involving cysteine (C), methionine (M), histidine (H) and tryptophan (W).

**Figure 1.**
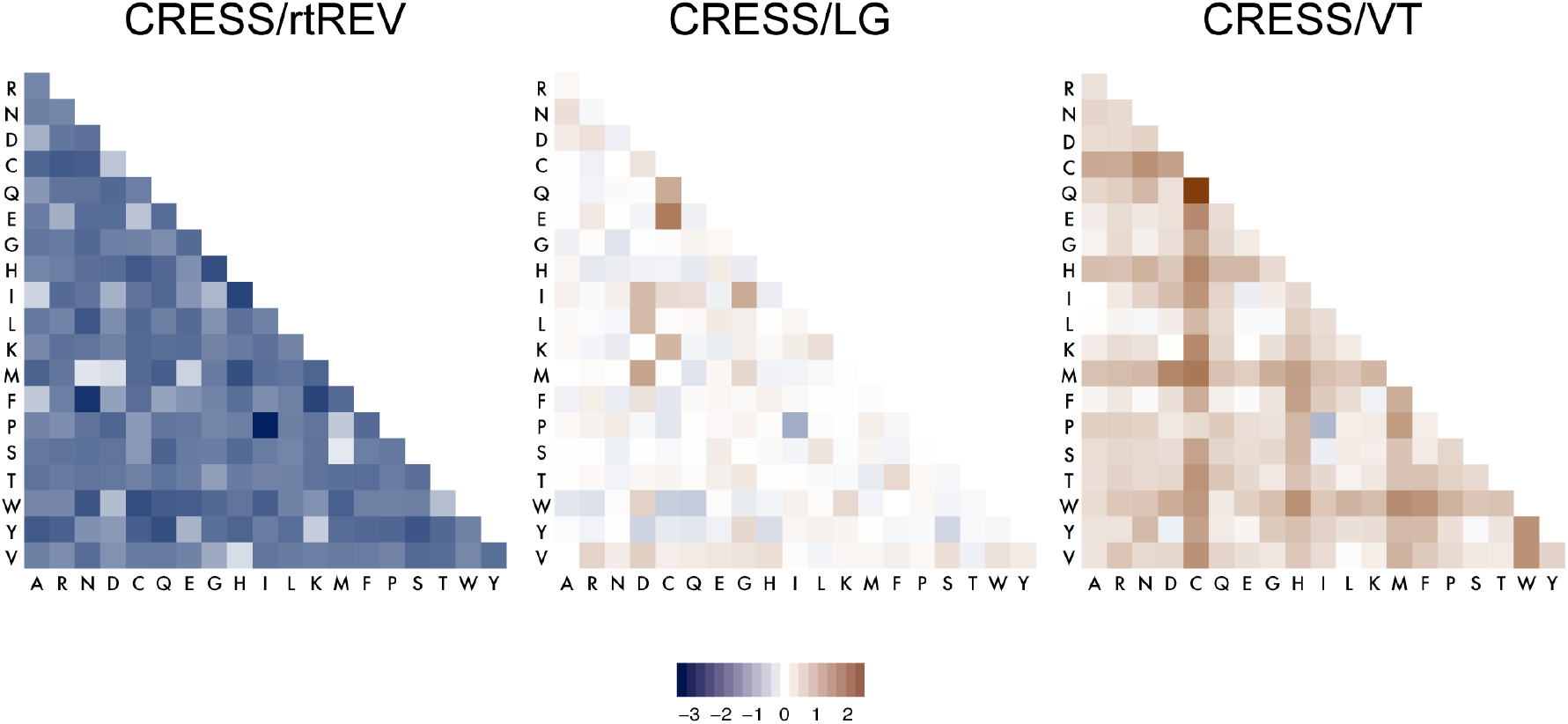
Comparison of four amino acid substitution matrices. Log-ratio rate matrices of CRESS rates over those of rtREV, LG and VT are shown left to right. The intensity of blue color indicates substitutions where CRESS has lower rates, and the intensity of brown color indicates substitutions where CRESS has higher rates.

### Genealogy construction and comparison

Four aLRT SH-like support maximum likelihood trees were built using CRESS, rtREV, LG and VT in PhyML3. 1000 bootstrap trees were also generated for each matrix in RaxML, and the bootstrap support were mapped to the PhyML trees (Figure 2 and Supplementary Files 5&6). The viral sequence-derived matrices (rtREV, CRESS) constructed more similar trees than those constructed with the two general matrices (LG, VT), which were more similar to each other (Figure 2, Supplementary File 6). Both virus-specific matrices placed *Nanoviridae* as a sister group to Alphasatellites (CRESS aLRT SH-like 0.942, rtREV aLRT SH-like 0.765, not supported by bootstrapping in either tree), while the general matrices placed *Nanoviridae* inside the Alphasatellite clade. There is a single alphasatellite sequence that groups with the geminiviruses; this is Ageratum leaf curl Cameroon alphasatellite, which appears to be a truncated version of the geminivirus Ageratum leaf curl Cameroon virus instead of sharing ancestry with other alphasatellites – its Rep belongs with other geminiviruses. The general matrices place *Smacoviridae* within clade containing *Geminiviridae*, *Genomoviridae* and some unclassified viruses, but the virus-specific matrices placed the *Smacoviridae* clade outside of clade with geminiviruses and genomoviruses.

**Figure 2.**
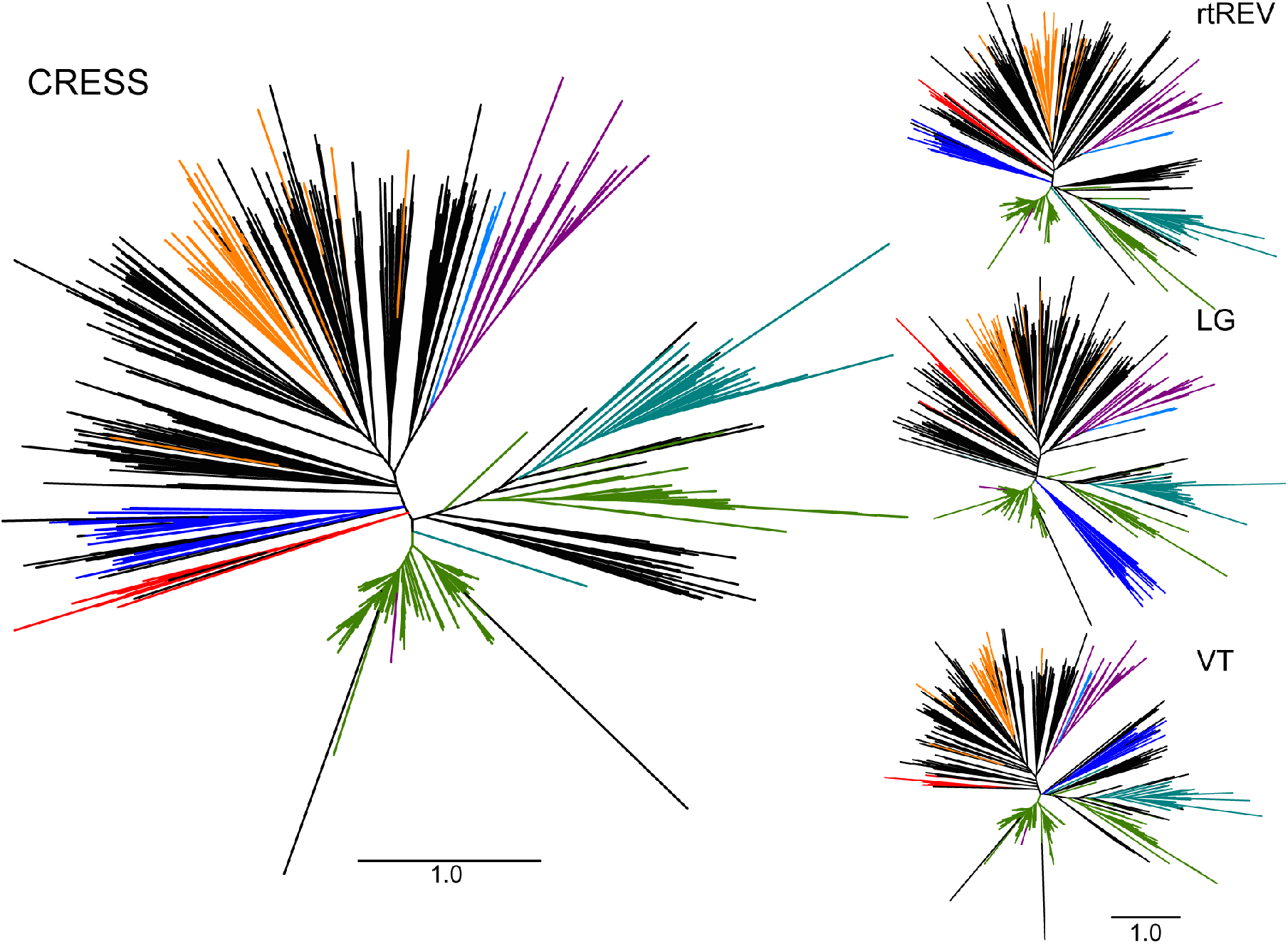
Left panel: Maximum likelihood tree of the Rep protein built using the CRESS matrix. Clockwise from the top: light blue for *Nanoviridae,* purple for the alphasatellites *(Alphasatellitidae),* teal for *Genomoviridae*, green for *Geminiviridae*, red for *Bacilladnaviridae*, dark blue for *Smacoviridae*, and orange for *Circoviridae*. Black taxa are from currently unclassified CRESS DNA viral sequences. The scale bar shows branch length of 1.0 substitution per site. A version of this tree with accession number labels at the tips for all taxa can be found in Supplementary File 5. Right panel: Maximum likelihood trees of the Rep protein built using the rtREV, LG and VT matrices. All three trees share the scale bar below, which shows branch length of 1.0 substitution per site. NEXUS files for these trees with aLRT support values can be found in Supplementary File 6.

The CRESS matrix tree is the only tree that grouped all currently classified *Bacilladnaviridae* in one clade (aLRT SH-like 0.833, not supported by bootstrapping) – the trees built with the other three matrices failed to place *Thalassionema nitzschioides* DNA virus Rep (BAN59850) inside the *Bacilladnaviridae* clade. And finally, classified *Circoviridae* are often intermingled with unclassified Rep sequences, as have seen in previous Rep trees (Kazlauskas et al., 2018; Kraberger et al., 2015a; Simmonds et al., 2017; Varsani and Krupovic, 2018).

The most surprising relationship resolved, which was found in all four trees, is the single clade for the intron-containing form of Rep from two CRESS DNA virus families. The CRESS matrix tree has 55% bootstrap support and 0.997 aLRT SH-like support for the clade containing all but one member of *Genomoviridae* and the intron-containing geminivirus Reps (from the genera *Becurtovirus, Capulavirus, Grablovirus* and *Mastrevirus*), while the three other matrices produced 45% bootstrap support, and 0.915-0.997 aLRT SH-like support for the clade (Figure 2 and Supplementary Files 5&6). Further inspection of the unclassified Reps in this clade showed most (7/11) contained annotated introns in the same location in their GenBank files. Blastp results indicated the remaining four Reps are highly similar to intron-containing mastrevirus Reps, genomovirus Reps and other unclassified intron-containing Reps, implying that all members of this clade contain an intron that they inherited by descent. We investigated the support in our alignment for the single ancestral origin of intron-containing Reps in these two families by dividing the MUSCLE aligned full dataset into helicase and endonuclease domains as described in previous literature (Kazlauskas et al., 2018). The two resulting trees independently show support for the intron-containing form of Reps (endonuclease tree aLRT SH-like support 0.842, helicase tree aLRT SH-like support 0.909, (Figure 3; Supplementary Files 7 and 8)).

**Figure 3.**
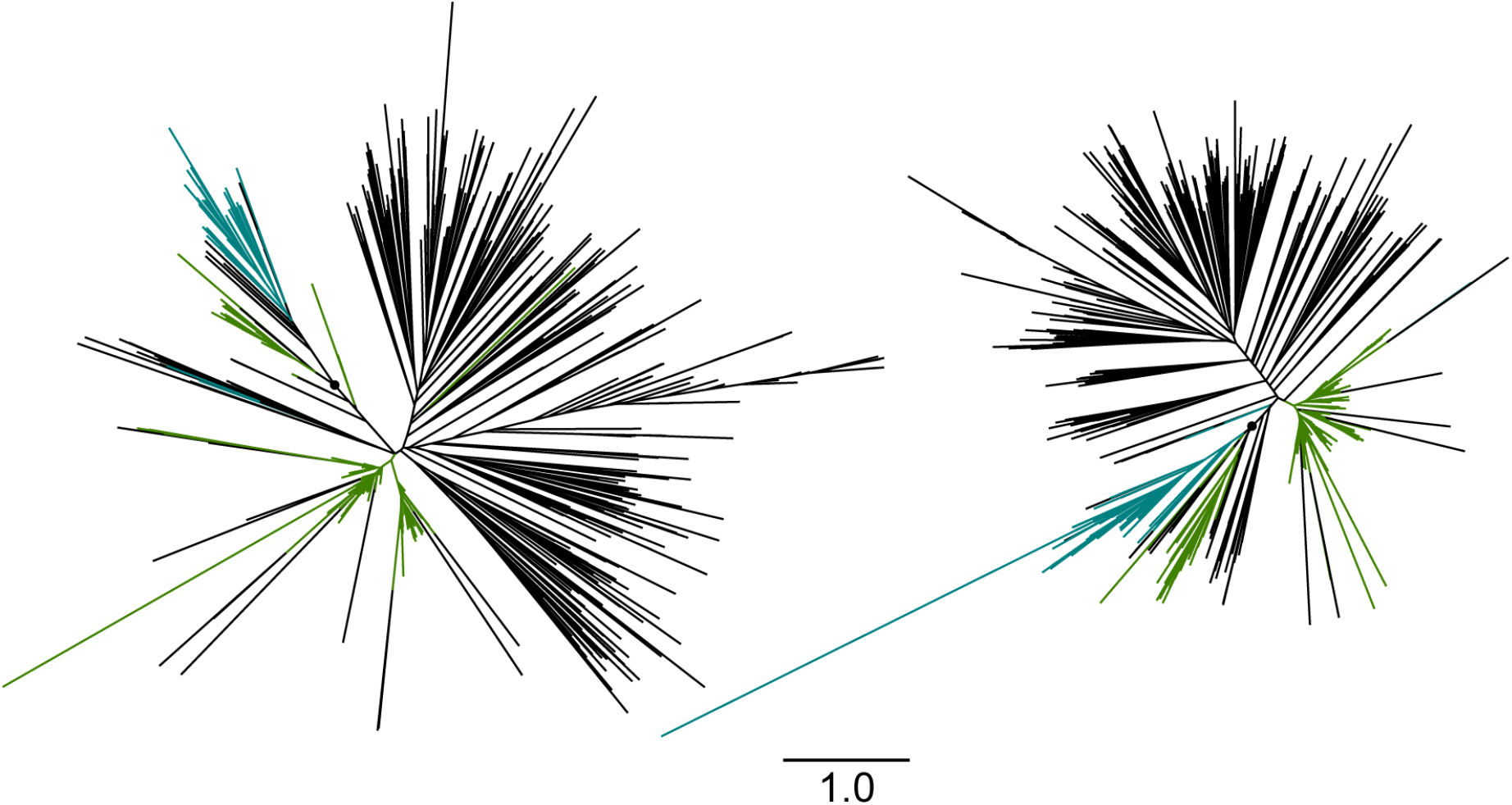
Unrooted maximum likelihood trees of CRESS Rep helicase (left) and endonuclease (right) domain. Genomovirus Reps are colored in teal, geminivirus Reps are colored in green, all other taxa (classified and unclassified) are colored in black. The black circles indicate the nodes for the common ancestor of intro-containing Reps (helicase tree: aLRT SH-like support 0.909; endonuclease tree: aLRT SH-like support 0.842). A version of these trees with accession number labels at the tips for all taxa can be found in Supplementary Files 7 and 8.

### Single origin for intron-containing Rep

The single origin of intron-containing Reps from *Genomoviridae* and four genera within *Geminiviridae*, from an unspliced Rep is a result not previously observed in the literature, which repeatedly show Reps of *Genomoviridae* and *Geminiviridae* as reciprocally monophyletic (e.g., (Dayaram et al., 2015a; Simmonds et al., 2017; Zawar-Reza et al., 2014). The grouping of intron-containing Reps is appealing from the perspective of Occam’s razor: it is more plausible that an intron was only inserted once in the same location in evolutionary history of the CRESS DNA viruses instead of being inserted at the same location multiple times in two separate lineages. We investigated why others have not observed this single intron-containing clade in previous trees. Our analysis included RefSeq sequences for each CRESS DNA viral species, including from the *Geminiviridae,* which is the most speciose viral family, while most researchers use only a small number of representative geminiviruses in a dataset, and these representatives most often come from *Begomovirus,* which comprises 75% of the annotated geminivirus species (https://talk.ictvonline.org/taxonomy/). Begomovirus Reps do not contain an intron, so often the sequence diversity of geminivirus Reps was inadequately represented (Castrignano et al., 2017; Simmonds et al., 2017; Varsani and Krupovic, 2018). Interestingly, when we reduced the number of *Begomovirus* Reps in our dataset to the same size as the largest intron-containing genus, *Mastrevirus* (n=37), the single origin of intron-containing geminivirus and genomovirus Reps loses support, and nanoviruses group within the alpha-satellites instead of being reciprocally monophyletic (Figure 4). We also investigated whether our trimming of our alignment affected our results: trimmed and untrimmed alignments of our 926 sequences did not change major tree branching patterns including the monophyly of intron-containing Rep sequences (data not shown).

**Figure 4.**
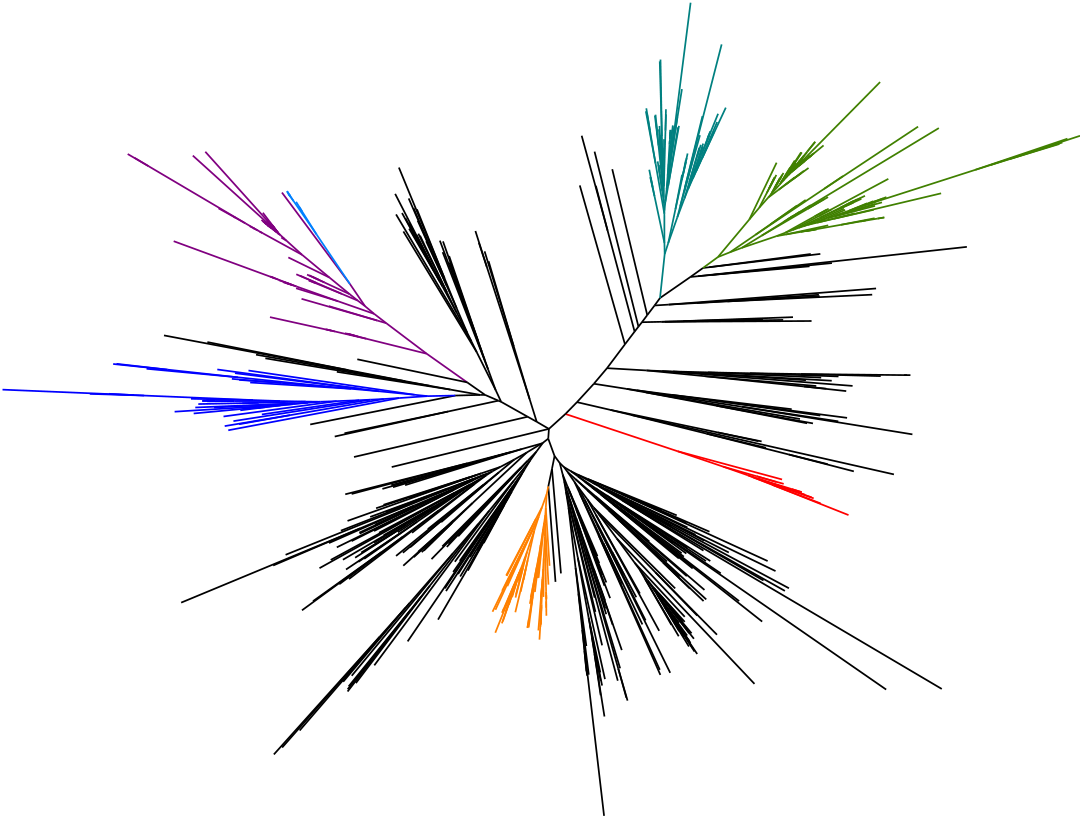
Unrooted maximum likelihood trees with equal number of *Begomovirus* and *Mastrevirus* Reps built with CRESS matrix. Alignment produced by MUSCLE. Green for *Geminiviridae,* teal for *Genomoviridae*, dark blue for *Smacoviridae*, red for *Bacilladnaviridae*, orange for *Circoviridae*, light blue for *Nanoviridae*, purple for the alphasatellites (*Alphasatellitidae*). Currently unclassified taxa are colored black. A version of this tree with accession number labels at the tips for all taxa can be found in Supplementary File 9.

To further probe the difference between our tree and a recently published Rep tree that showed members of *Geminiviridae* and *Genomoviridae* as separate clades (Kazlauskas et al., 2018), we examined the methods used to make the two Rep trees. The previously published study used a smaller dataset of CRESS DNA virus Rep sequences (n=647), which was further pruned by removing detectable recombinants (final n = 380). When we removed the recombinant sequences detected in that study from our dataset, we still supported the single origin of intron-containing Reps in *Genomoviridae* and the appropriate genera from *Geminiviridae* (Figure 5). Additionally, there were differences in alignment algorithms, as the other study used MAFFT instead of MUSCLE. We aligned our original dataset with MAFFT and compared its ML tree (Figure 6) with our original MUSCLE alignment derived tree (Figure 2). The MAFFT tree of our entire dataset did not show single intron origin for intron-containing members of *Geminiviridae* and *Genomoviridae.* However, when we extracted the geminivirus and genomovirus sequences from both the MAFFT alignment and the original MUSCLE alignment and built smaller trees (Figure 7), both showed the intron-containing clade of geminiviruses and genomoviruses separate from the non-intron-containing clade of geminiviruses. This suggests that the inclusion of Reps from other families obscured the potential single origin of intron-containing Rep in the MAFFT alignment tree. While both algorithms have been used to analyze CRESS DNA virus evolution, there is no *a priori* or *a posteriori* reason to prefer one algorithm over the other. Regardless, some datasets aligned with both algorithms provided support for a single origin of intron-containing Reps in families *Geminiviridae* and *Genomoviridae*.

**Figure 5.**
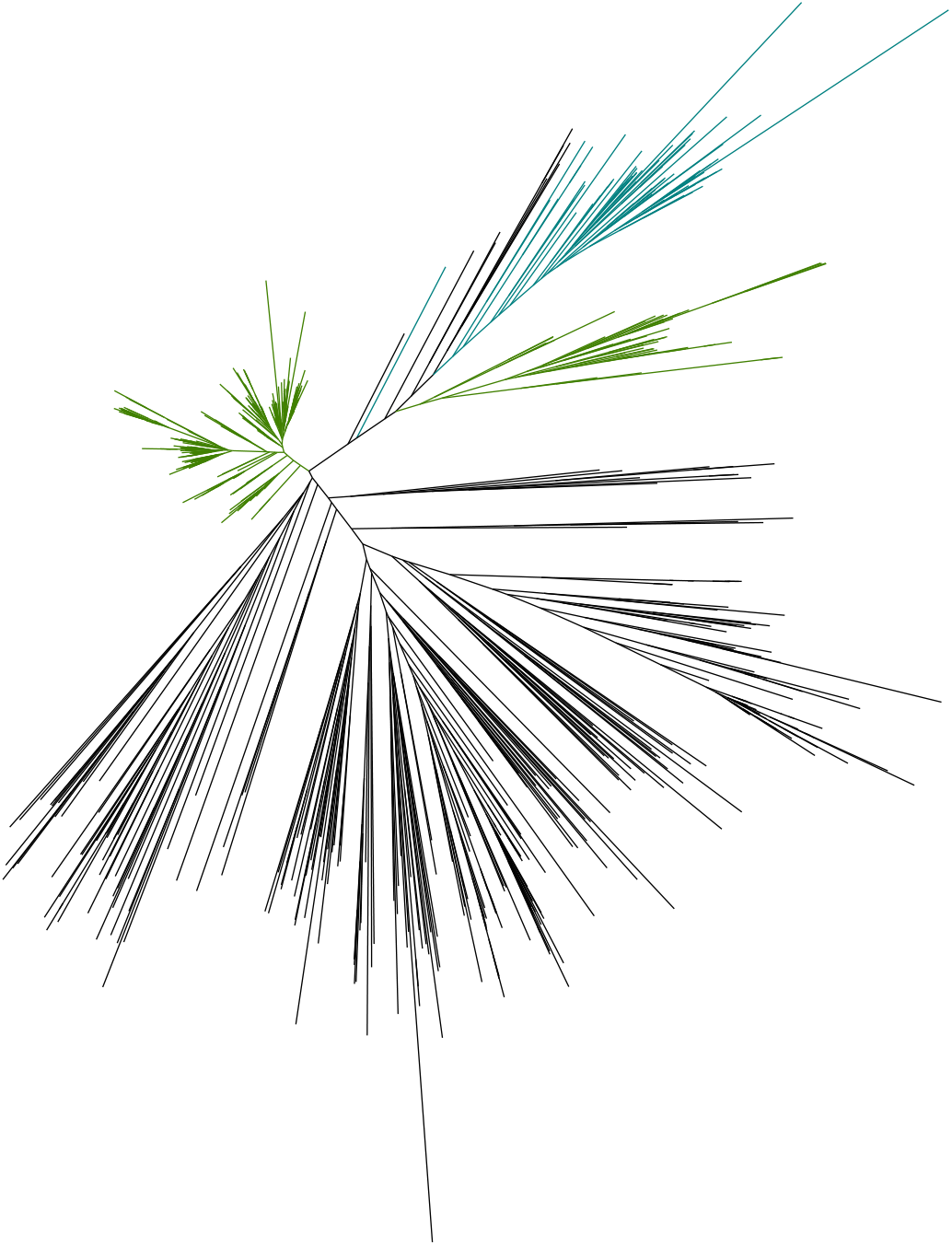
Unrooted maximum likelihood tree built with 123 recombinant sequences removed from the 926 CRESS Rep dataset. Green for *Geminiviridae,* teal for *Genomoviridae,* all other taxa (classified and unclassified) are black. A version of this tree with accession number labels at the tips for all taxa can be found in Supplementary File 10.

**Figure 6.**
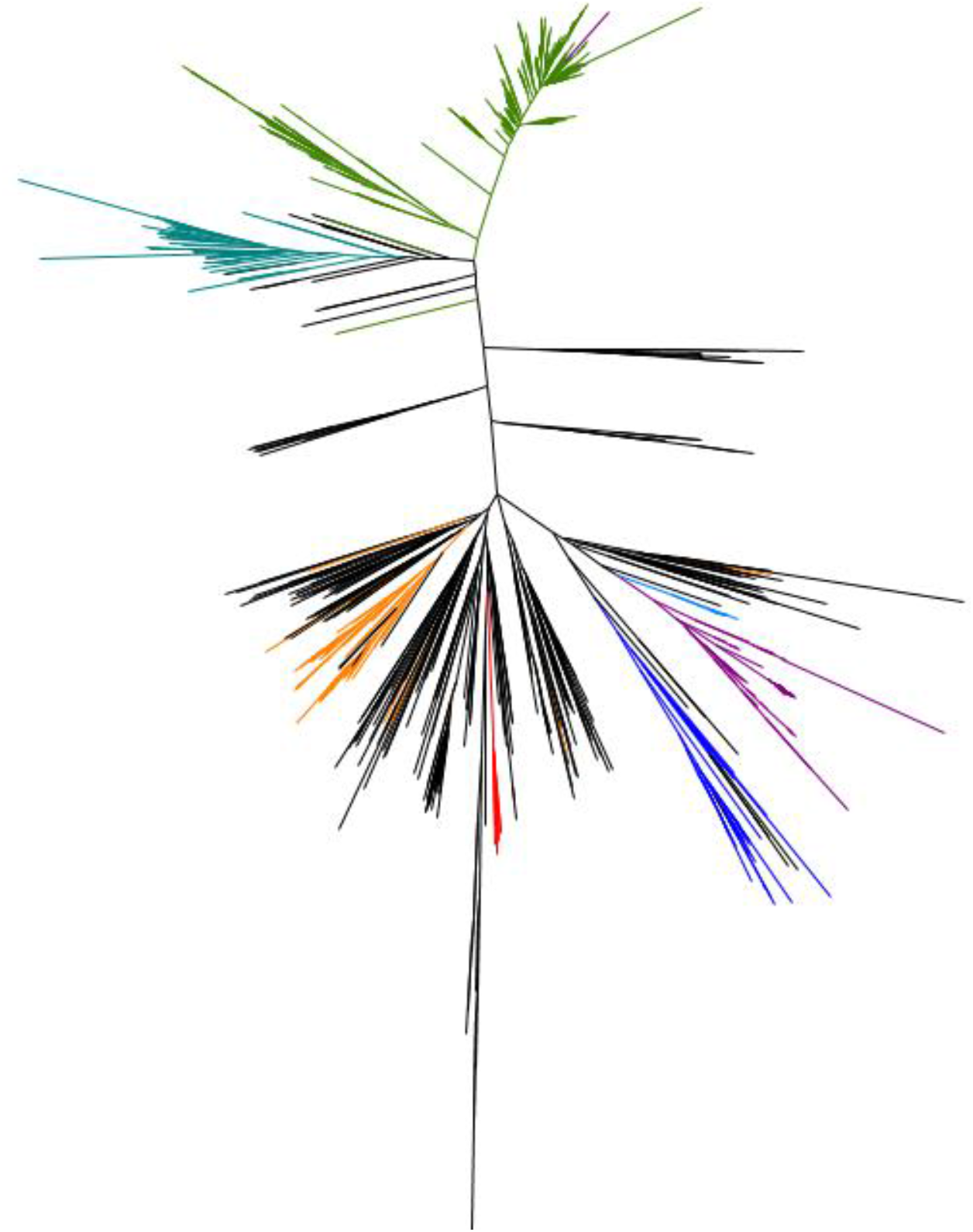
Unrooted maximum likelihood tree built with the full 926 CRESS Rep dataset. Alignment produced by MAFFT. Green for *Geminiviridae*, teal for *Genomoviridae*, dark blue for *Smacoviridae*, red for *Bacilladnaviridae*, orange for *Circoviridae*, light blue for *Nanoviridae*, purple for the alphasatellites (*Alphasatellitidae*). Currently unclassified taxa are black. A version of this tree with accession number labels at the tips for all taxa can be found in Supplementary File 11.

**Figure 7.**
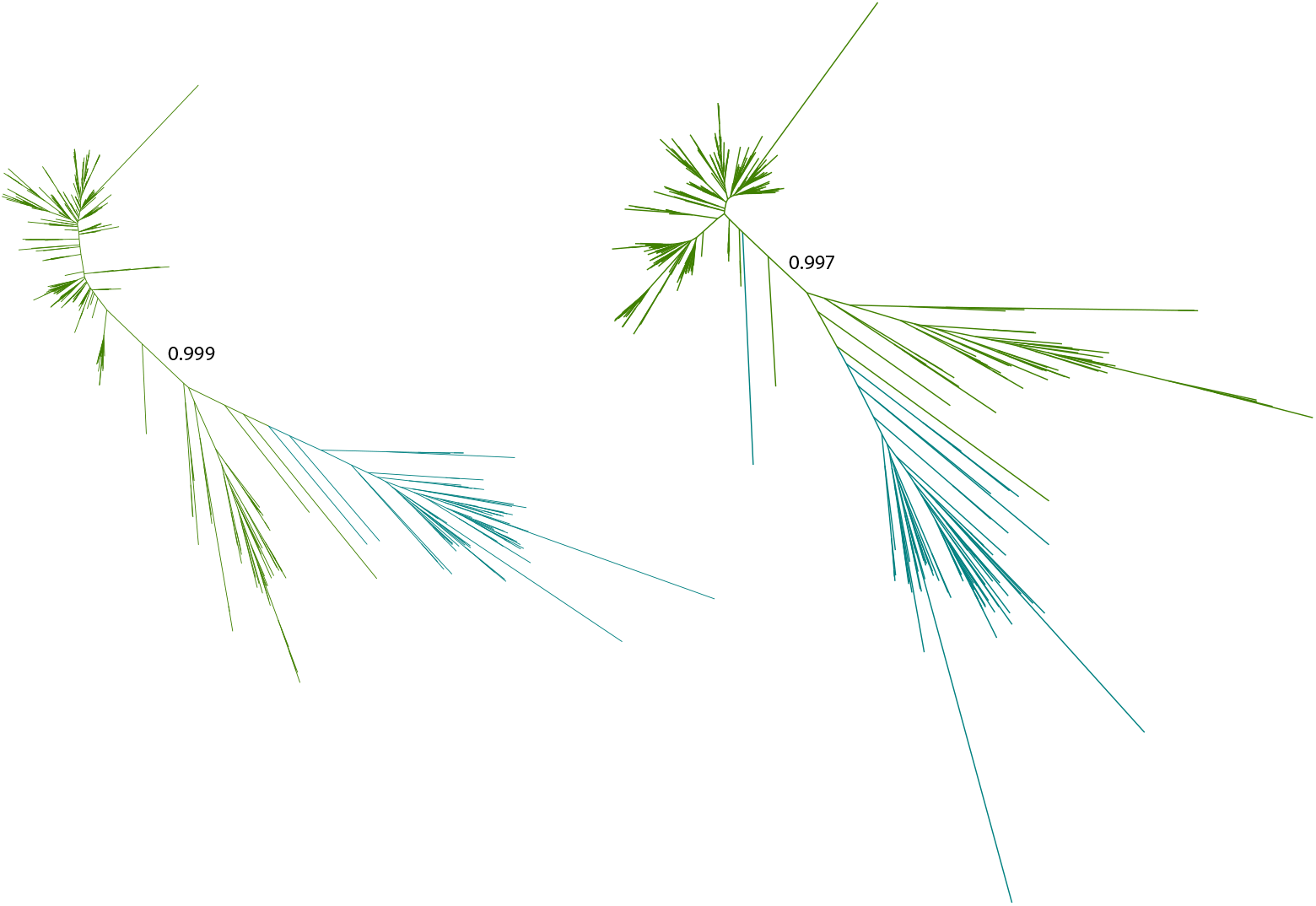
Unrooted maximum likelihood tree with Geminivirus and Genomovirus Reps built with CRESS matrix. Alignment produced by MAFFT (left) and MUSCLE (right). Green for *Geminiviridae,* teal for *Genomoviridae.* SH-like support labeled on the branch between the no intron Reps (above) and intron-containing Reps (below). Versions of these trees with accession number labels at the tips for all taxa can be found in Supplementary Files 12 and 13.

### CRESS matrix performance on other sequences

Capsid protein (CP) sequences from CRESS DNA viral families *Bacilladnaviridae, Circoviridae, Genomoviridae*, *Nanoviridae* and *Smacoviridae* were downloaded and aligned using MUSCLE. Model performance was ranked in ProtTest3, comparing 56 different models (CRESS, LG, VT, rtREV, HIV-B, HIV-W, FLU in combination with +I, +G, +F parameters). The CRESS models (CRESS+G+F, CRESS+I+G+F) outperformed all other models in building ML trees with CRESS DNA viral CP alignments (Table 1). An alignment of NS1/Rep sequences from linear ssDNA parvoviruses was also tested, CRESS+I+G+F was the best model chosen based on three different criteria (Table 1). A circular ssDNA phage major capsid protein multiple sequence alignment was provided by collaborators (K. Rosario and M. Breitbart, details in (Creasy et al., 2018) and LG+I+G+F outperformed all other matrices tested, though the CRESS matrix was ranked second. These results indicate that the CRESS matrix can describe both Rep and CP evolution of eukaryote-associated ssDNA viruses, and potentially would be useful for some datasets of ssDNA phage protein sequences in the future.

**Table 1.**
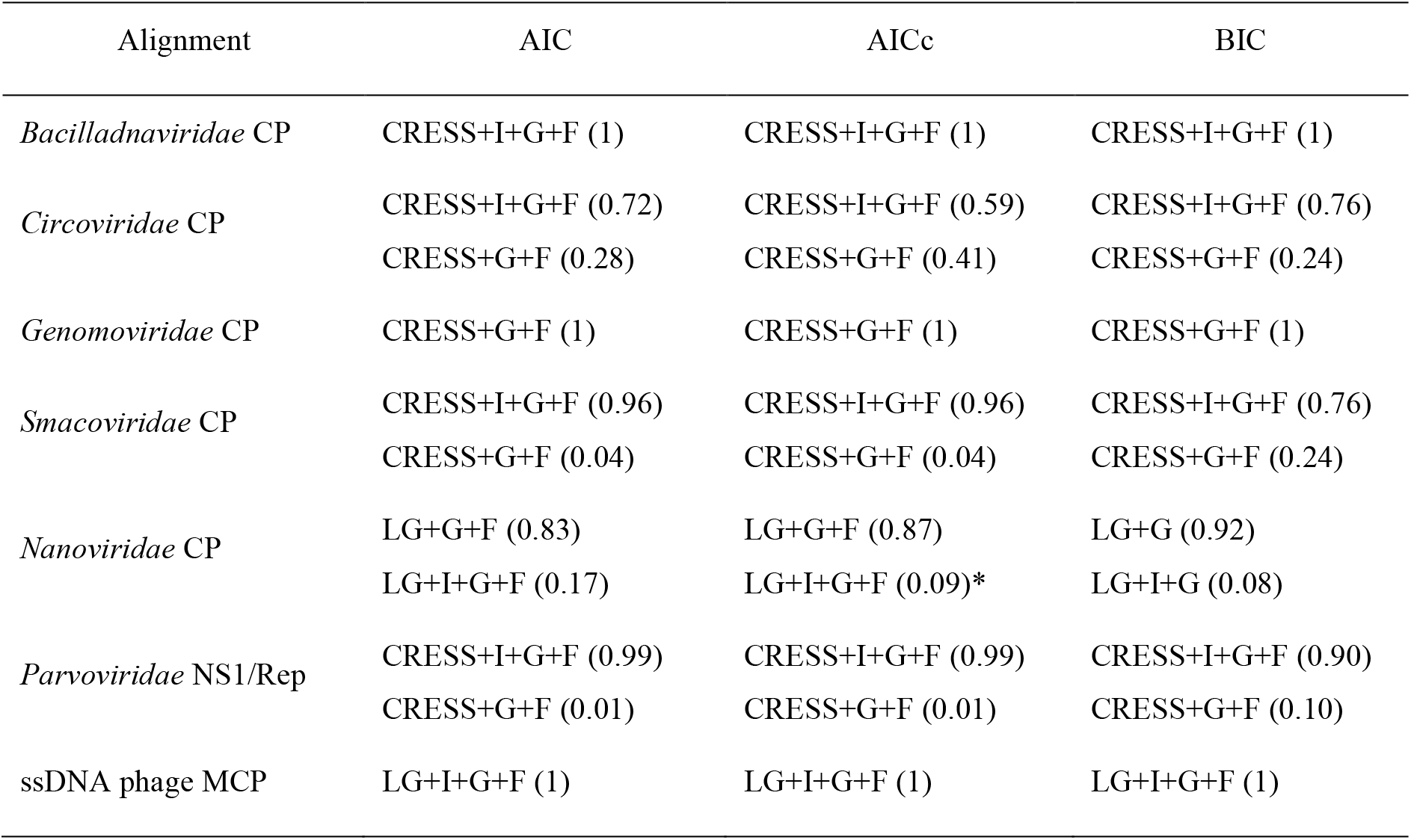
Best model chosen by ProtTest3 for each tested ssDNA protein alignment. The table shows the models chosen by these information criteria and their corresponding weights. The best performing models were ranked by Akaike information criterion (AIC), corrected Akaike information criterion (AICc) and Bayesian information criterion (BIC) scores and the weights of these models are shown in parenthesis. +G assumes gamma-distributed rate variation across sites. +I estimates the proportion of invariant sites, +F uses empirical amino acid frequency. * indicates that the top two models’ weights do not sum to 1.

## Discussion

In this study, we built a CRESS DNA viral Rep genealogy with a bespoke CRESS matrix to gain insight into the relationships among the diverse families of CRESS DNA viruses. We observed an unprecedented single evolutionary origin for intron-containing Rep shared by some members of the *Geminiviridae* and virtually all in *Genomoviridae.* In fact, the one genomovirus Rep (from KY056250) that was not in this clade is likely inaccurately classified – it was named a gemycircularvirus due to its 52% similarity in the CP to a gemycircularvirus sequence obtained from preserved caribou feces (Lima et al 2017 JGV). Its predicted Rep does not appear to have an intron and is unlike the Reps of other characterized genomoviruses, so it is appropriately outside of the supported clade. Further supporting its misclassification is that the conserved nonanucleotide in the genomic origin of replication (TAAGATTCG, Lima et al 2017) does not match that of genomoviruses (TAWWDHWAN, Varsani and Krupovic 2017) or any family of CRESS DNA viruses. We propose that this sequence should not be considered a member of *Genomoviridae* and that genomovirus Reps form a well-supported clade nested within a well-supported monophyletic group of intron-containing Reps. It should be noted that some members of *Circoviridae* are also predicted to have an intron in their Rep gene; this is not in the same location as in geminiviruses and genomoviruses, and likely represents a unique integration event (Mankertz and Hillenbrand, 2001). Intron splicing in the circoviruses has only been studied in the porcine circoviruses, which form a well-supported clade (aLRT SH-like 0.994, 100% bootstrap, Figure 2) – our Rep genealogy also supports the monophyly of this second intron introduction into Rep.

### Relationships among other CRESS DNA viral families

Some of the large branching patterns observed on our full dataset tree are consistent with previous analyses. Notably, the close association of members of *Nanoviridae* and alphasatellites, and the close placement of members of *Genomoviridae* and *Geminiviridae*. In contrast, without strong support, bacilladnavirus and smacovirus Reps were close to each other in our tree compared to previous trees (Dayaram et al., 2015b; Varsani and Krupovic, 2018). The relatively few members of *Bacilladnaviridae* are known to infect diatoms (Kazlauskas et al., 2017; Nagasaki, 2008), which are phylogenetically closer to plants than animals. Therefore, we might expect their Reps to be more similar to other plant-infecting CRESS DNA viruses *(Nanoviridae, Geminiviridae)* instead of smacoviruses, which are associated with animals. In previous trees, Reps from *Smacoviridae* closely grouped with those of nanoviruses and alphasatellites, with strong support (Kazlauskas et al., 2018).

A majority of the Reps from unclassified CRESS DNA viruses are sister taxa to circovirus Reps. *Circoviridae* used to be the catch-all group for all circular eukaryotic infecting viruses not associated with plants, and species assigned to *Circoviridae* but not to a genus may represent additional genera or families of CRESS DNA viruses (Rosario et al., 2017). As previously proposed, many of the unclassified CRESS DNA viruses should be organized into novel families (Kazlauskas et al., 2018).

Some unclassified Reps were placed in unexpected places in the CRESS matrix tree, such as the two protruding, unclassified taxa inside the Geminivirus clade. This suggests that the dataset does not have many sequences that are similar to these unclassified Reps, so the alignment algorithm was not able to accurately align this sequence to other sequences present in the dataset. Perhaps these long branches are due to poorly aligned Rep sequences, which could make their placement near geminiviruses coincidental; perhaps they are the first representatives of highly derived geminivirus-like groups. Updated annotation is pending for many CRESS DNA viral sequences in GenBank, both currently classified and unclassified, so the odd placement of some potentially mislabeled Rep sequences may be more understandable in the future.

While we do not know what the true phylogeny of CRESS DNA viruses looks like, especially as discovery of this group is still ongoing, we believe that our comprehensive genealogy is the best representation of the relationship among the diverse Rep sequences that have been sampled to date. The bespoke CRESS matrix will help future phylogenetic analyses on CRESS DNA viruses as others discover additional, novel representatives of this group.

### The utility of the CRESS matrix

Prior to phylogenetic reconstruction, tools such as JModelTest and ProtTest are widely used to select for the most appropriate model with which to estimate phylogenetic relationships for a given alignment. General models such as Dayhoff, BLOSUM, VT, WAG, LG, are all estimated from unconstrained, nonspecific, aligned protein sequences. The most recent general model, LG, published in 2008, used the entire Pfam database (Finn et al., 2016), but the database is heavily weighted towards non-viral sequences (Skewes-Cox et al., 2014). Unfortunately, the majority of CRESS DNA viruses were discovered and accessioned into NCBI GenBank after 2009 (Zhao et al., 2019), which means any genomic novelty in this group would be excluded from the LG matrix. Furthermore, since most CRESS DNA viruses have very few ORFs, often only a CP and a Rep, their sequence diversity contributes little to the overall Pfam database. While general models do increasingly well at describing the patterns dominated by genetic code constraints and the physiochemical properties of the amino acids (Murrell et al., 2011), there will always be opportunities for specific matrices to resolve protein evolution in biological entities that have more unique lifestyles and constraints, such as fast-evolving CRESS DNA viruses with single-stranded mutational biases (Cardinale et al., 2013; Frederico et al., 1990; Xia and Yuen, 2005).

Among the four matrices used in this study, it seems the CRESS matrix falls in the large gulf between the fast-changing rtREV and the slower general matrices (Figure 2). While we have not compared CRESS to all described matrices, it may fill a useful niche for other proteins’ evolution. Just as the rtREV model has been useful beyond studies on retro-transcribing elements, the inclusion of the CRESS matrix in ProtTest may result in the model being used to develop phylogenies for proteins from distantly related viruses. Perhaps its moderate substitution rate would be preferred by other viruses that evolve faster than cells but more slowly than some RNA viruses (Duffy et al., 2008).

We found the fmg algorithm to perform better than fit. Importantly, fmg successfully extracted a consistent signal of protein evolution that overcame the stochastic effects of different randomly split datasets, regardless of whether LG or VT was used as the seed matrix. For the creation of future specific amino acid models, when known phylogenies are not available (cf. rtREV (Dimmic et al., 2002) and HIV (Nickle et al., 2007a)), our work recommends FastMG.

Our CRESS matrix was useful both for building our Rep genealogy and describing the evolution of other proteins in CRESS DNA viruses and linear ssDNA viruses. As the annotation of ssDNA phages known by sequences alone improves, we welcome the opportunity to test the fit of the CRESS matrix to their Rep homologs – and are curious if CRESS would outcompete established matrices to describe the evolution of phage Reps, which are very distantly related to eukaryote-associated Reps (Koonin and Ilyina 1993). Regardless, we anticipate the CRESS matrix will likely become the consistently chosen matrix for eukaryotic CRESS DNA virus protein phylogenies, and hopefully the substitution rates estimated from CRESS DNA viral sequences helps accurately model their protein evolution, creates more accurate alignments, and is useful in searching databases for similar sequences (Thorne, 2000).

## Material and Methods

### Dataset generation

Collection: Rep sequences were downloaded from NCBI RefSeq December, 2017. RefSeq sequences were chosen to include unclassified viral sequences but exclude repeating sequences from the same species. Some CRESS DNA viruses have introns in their Rep genes. Generally, the spliced protein product is specified in GenBank and that was used, however we had to manually form the spliced Rep from some older sequences of families with known introns (i.e., *Mastrevirus).* This was done after MUSCLE v3.8.31 (Edgar, 2004) alignment of all *Geminiviridae* Reps, and the unaligned region corresponding to the introns were cut. The list of included sequences with details of edits in Supplementary File 14 and the alignment is in Supplementary File 15.

Alignment and trimming: All Rep sequences were pooled into one FASTA file and aligned using MUSCLE v3.8.31 (default setting: max 16 iterations) (Edgar, 2004). The multiple sequence alignments were trimmed using trimAl v1.2 command line (Capella-Gutierrez et al., 2009), resulting in alignments with the length of 329 aa, the average length of the Rep data set. This was achieved by removing all columns with gaps in more than 40% (-gt 0.6) of the sequences while respecting the conservation of 17.12% of the columns (trimAl v1.2 adds columns in decreasing order of score when necessary). The trimmed MUSCLE alignment was used for matrix estimation.

### Matrix estimation

ProtTest 3.4 (Darriba et al., 2011) determined the best model for building the initial maximum likelihood (ML) tree of the trimmed MUSCLE alignment. A total of 80 models’ NNI ML trees were compared: JTT, LG, Dayhoff, WAG, Blosum62, VT, rtREV, DCMut, MtREV, MtArt with combinations of +I, +G, +F. VT+G+F was the overall best performing model according to AIC, AICc, BIC scores. A full data set ML tree was constructed with VT+G+F using PhyML 3.1(Guindon et al., 2010) for subsequent computations.

Then, the 926-sequence alignment was randomly jackknifed using a python script (https://github.com/lzhao-virevol/matrix) into two halves as the training and test datasets. Ten pairs of jackknifed datasets were generated as replicates. The initial maximum likelihood tree was also divided accordingly into ten training trees and ten test trees manually using Figtree (http://tree.bio.ed.ac.uk/software/figtree/). The training set was used to estimate an amino acid substitution matrix with a modified HyPhy batch file from Nickle et al. (Nickle et al., 2007a) (https://github.com/lzhao-virevol/matrix) and FastMG (Dang et al., 2014). We used two matrices to seed our matrix estimation: VT, the best-fitting matrix to the full dataset and LG, the most recent general amino acid substitution matrix (Le and Gascuel, 2008), which is frequently used to describe CRESS DNA virus evolution (Bistolas et al., 2017; Castrignano et al., 2017; Kaszab et al., 2018; Male et al., 2016). The HyPhy estimated matrices initiated with LG and VT are named fitLG and fitVT matrices. The FastMG estimated matrices initiated with LG and VT are named fmgLG and fmgVT matrices.

### Matrix evaluation

Likelihood scores were calculated for each matrix of interest (LG, RtREV, VT, fitLG1-10, fitVT1-10, fmgLG1-10, and fmgVT1-10) describing the ten test datasets (half alignment and tree) in PAML: codeml using model 3, Empirical+F (Yang, 1997). We included the LG and VT matrices that the estimated matrices were based on, and included rtREV because it is a specific matrix that has been previously used to describe CRESS DNA virus Rep evolution (Dayaram et al., 2016; Dayaram et al., 2015a; Dayaram et al., 2015b; Kraberger et al., 2015b; Rosario et al., 2015). To examine how similar the estimated matrices are to each other (and to the three established matrices), we calculated Pearson correlations in Excel (Redmond, WA) for the matrices.

The best-fitting matrix for of the ten test sets according to codeml results was used to build a maximum likelihood tree (PhyML3 with 4 discrete gamma rate categories, and empirical amino acid frequency options), producing ten test sets trees each. The maximum likelihood scores of these 100 trees were rank ordered by-In score within each test set. The matrix with the highest overall ranking was chosen as the best performing matrix, named the CRESS matrix. We compared the CRESS matrix to the established matrices VT, LG and rtREV using log10 ratios (Excel).

### Dataset size validation

A series of datasets with identical sequence number and length as the training datasets were simulated to test if the training sets are sufficient to recover the amino acid substitution patterns. These pseudo datasets were simulated under the WAG model (Whelan and Goldman, 2001), with different site rate variation parameters (alpha: 0.1, 0.5, 1, and 2) in Seq-Gen v1.3.3 (Rambaut and Grassly, 1997). We simulated alignments of similar size (463 sequences with the length of 329aa) to the training set Rep alignment.

Four alignments (one for each parameter for the gamma distribution) were combined with the tree from training set 10 to produce eight substitution matrices by the same matrix estimation methods used to generate the CRESS Rep-derived matrices: four fmg and four fit (Dang et al., 2014; Nickle et al., 2007a), both using seed matrix VT. The sequences-specific amino acid substitution matrix of each pseudo-dataset was estimated using the algorithms described above, and then compared to WAG through Pearson correlation tests.

### CRESS DNA viral Rep tree construction

Four aLRT SH-like supported maximum likelihood Rep trees were also built in PhyML3 using CRESS, VT, LG and rtREV matrices with +G+F options. 1000 bootstrapped trees were also estimated using RAxML v8.2.10 (Stamatakis, 2014) on CIPRES (RAxML-HPC2 on XSEDE) with +G+F options using CRESS, VT, LG and rtREV matrices. The bootstrapped trees were mapped onto corresponding aLRT SH-like supported ML trees with local non-MPI RAxML-HPC v8.1.17 (Stamatakis, 2014).

As there is a hotspot of recombination between the two domains of Rep (Kazlauskas et al., 2018), we separated the endonuclease and helicase domain alignments by splitting the trimmed MUSCLE aligned full-length Rep alignment, according to previously published descriptions (Kazlauskas et al., 2018). The endonuclease and helicase domain maximum likelihood trees were built in PhyML3 (CRESS+G+F).

We also removed sequences from our alignment that were determined to be recombinant by another study (Kazlauskas et al., 2018), and reduced the number of sequences used from the most overrepresented genus, *Begomovirus*. A total of 123 recombinants and 319 begomoviruses were separately removed from the full dataset’s MUSCLE alignment. The two trees were built in PhyML3 using CRESS+G+F.

We aligned the 926 Rep sequences using MAFFT v7. 271 (options L-INS-I-ep 0.123) (Katoh and Standley, 2013) and also extracted the taxa from the MAFFT alignment (and from the analogous MUSCLE alignment) that were members of *Geminiviridae* and *Genomoviridae* for additional analysis. Trees based on these three datasets were built using PhyML3 with CRESS matrix and +G+F options.

All trees were visualized with Figtree (http://tree.bio.ed.ac.uk/software/figtree/) and edited in Adobe Illustrator.

### Model comparison using modified ProtTest

Capsid protein (CP) sequences from several families of CRESS DNA viruses and *Parvoviridae* RefSeq NS1 sequences were procured from NCBI in May 2018. Lists of CRESS DNA viral genome accession numbers were downloaded from the relevant family description from the International Committee on the Taxonomy of Viruses (ICTV, talk.ictvonline.org/taxonomy/) and if a CP was not identified in the sequence then the CP ORF was predicted using the NCBI ORF caller and validated through BLAST (accession numbers in Supplementary File 16). The 102 RefSeq *Parvoviridae* NS1/Rep sequences were either confirmed through database label or through BLAST (Supplement File 1). Each CP dataset and the parvovirus NS1/Rep dataset were aligned using MUSCLE (default setting: max 16 iterations) and left untrimmed. The phage major capsid protein sequence alignment was from published dataset (Creasy et al., 2018), generously provided by K. Rosario (University of South Florida), and trimmed using-gappyout option in TrimAl 1.2. These alignments were provided to a modified version of ProtTest that includes the CRESS model (https://github.com/lzhao-virevol/matrix) to evaluate various model performance. All viral specific models (CRESS, rtREV, HIVb, HIVw, FLU) and two relevant general models (LG and VT) with different combinations of +G, +F, +I (a total of 56 models) were tested using strategymode: NNI maximum likelihood tree.

## Supporting information

Supplemental Files 1-16 (excepting 6 & 15)

Supplemental File 6, trees

Supplemental File 15, alignment

## Acknowledgements

This work was supported by the National Science Foundation’s Assembling the Tree of Life program, DEB1240049. We thank David Swofford for advice on data sufficiency validation methods, Li Li for developing the initial matrix estimation scheme, Steen Hoyer for technical help and support, and LaShanda Williams and Natasia Jacko for useful comments and support.

