## Supplemental Files 1-16 (excepting 6 & 15) for "A comprehensive genealogy of the replication associated protein of CRESS DNA viruses reveals a single origin of intron-containing Rep"

Supplementary File 1. Maximum likelihood scores of trees built by best matrices from 10 test set alignments.

| | | Test Set | | | | | | | | | |
|---|---|---|---|---|---|---|---|---|---|---|---|
| | | 1 | 2 | 3 | 4 | 5 | 6 | 7 | 8 | 9 | 10 |
| Best matrix | fmgVT-1 | -163729 | -166092 | -170039 | -163298 | -161822 | -163795 | -163296 | -163092 | -165413 | -159692 |
| | fmgVT-2 | -163649 | -166082 | -170040 | -163291 | -161776 | -163754 | -163313 | -163030 | -165399 | -159710 |
| | fmgLG-3 | -163660 | -166018 | -170105 | -163276 | -161817 | -163773 | -163346 | -163052 | -165400 | -159725 |
| | fitVT-4 | -163667 | -166037 | -169988 | -163324 | -161761 | -163739 | -163325 | -163021 | -165413 | -159685 |
| | fmgLG-5 | -163647 | -166037 | -170026 | -163282 | -161860 | -163775 | -163340 | -163045 | -165361 | -159727 |
| | fmgVT-6 | -163612 | -166027 | -169959 | -163315 | -161793 | -163836 | -163301 | -163058 | -165378 | -159695 |
| | fmgVT-7 | -163631 | -166013 | -169952 | -163263 | -161791 | -163755 | -163376 | -163082 | -165380 | -159657 |
| | fmgVT-8 | -163676 | -166050 | -170006 | -163280 | -161788 | -163754 | -163334 | -163131 | -165412 | -159745 |
| | fmgLG-9 | -163698 | -166062 | -170021 | -163291 | -161832 | -163771 | -163329 | -163038 | -165483 | -159734 |
| | fmgVT-10 | -163600 | -166014 | -169998 | -163254 | -161785 | -163735 | -163299 | -163041 | -165375 | -159761 |

Supplementary File 2. Pairwise Pearson correlation of amino acid substitution rates of matrices. The ten FastMG estimated matrices for each of the ten training datasets seeded with VT are named fmgVT, those seeded with LG are fmgLG; the ten HyPhy-estimated matrices are named fitVT and fitLG.

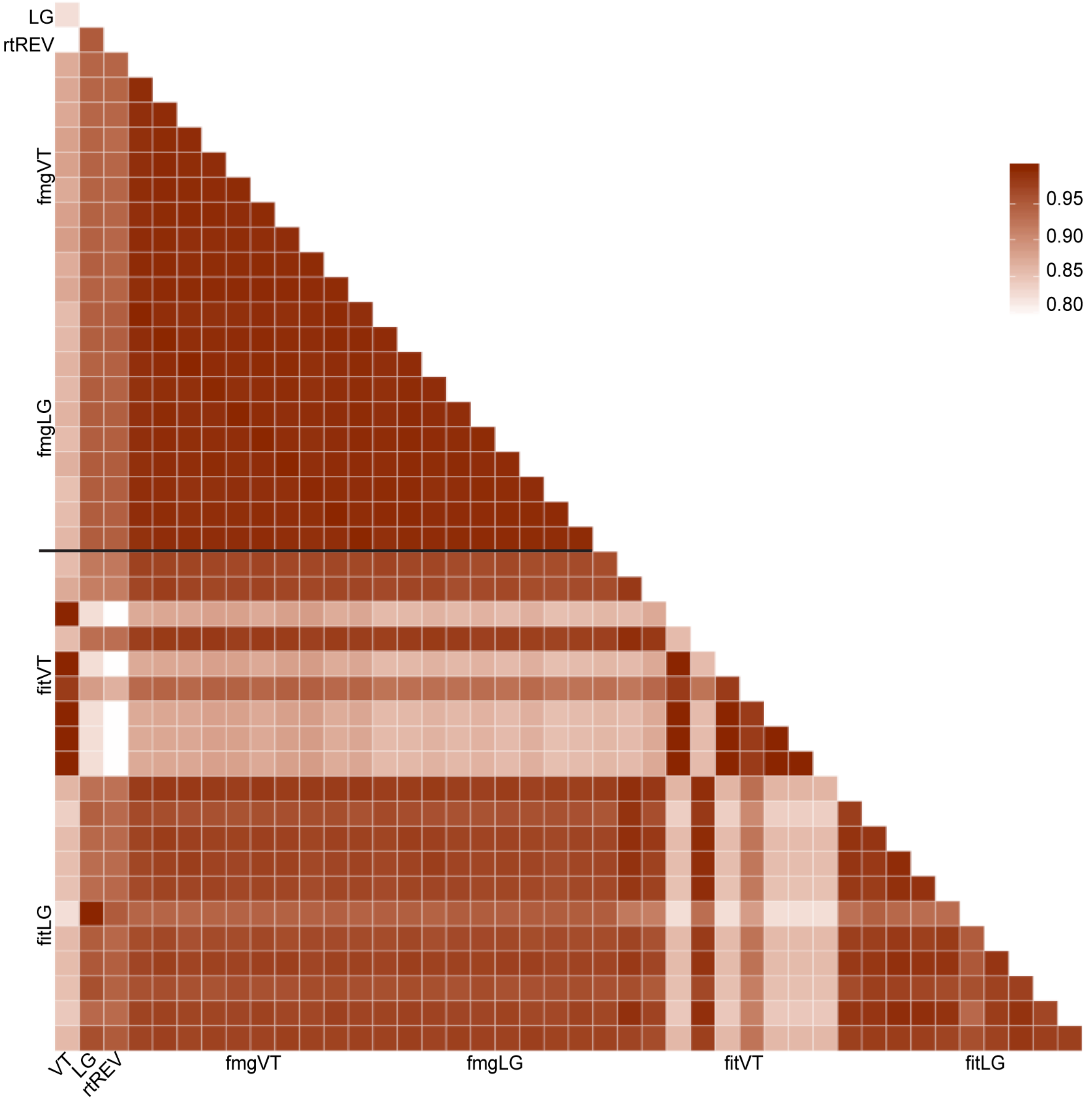

Supplementary File 3. Pearson correlation of log ratio of WAG rates over fmg method (top) and fit method (bottom) simulated data derived matrix rates. G01 means coefficient alpha for gamma distribution is 0.1 during simulation, G05 means coefficient alpha for gamma distribution is 0.5, G1 means coefficient alpha for gamma distribution is 1, G2 means coefficient alpha for gamma distribution is 2.

| | | | |
|---|---|---|---|
| WAG/fmg-G05 | 0.09 | | |
| WAG/fmg-G1 | 0.257 | -0.046 | |
| WAG/fmg-G2 | 0.263 | 0.33 | -0.02 |
| | WAG/fmg-G01 | WAG/fmg-G05 | WAG/fmg-G1 |

| | | | |
|---|---|---|---|
| WAG/fit-G05 | 0.049 | | |
| WAG/fit-G1 | 0.066 | 0.038 | |
| WAG/fit-G2 | 0.011 | 0.27 | -0.165 |
| | WAG/fit-G01 | WAG/fit-G05 | WAG/fit-G1 |

Supplementary File 4. Rank order of maximum likelihood scores of trees built by best matrices from 10 test set alignments. Best matrix with the lowest ranking is highlighted in peach color.

| | | test set | | | | | | | | | | |
|---|---|---|---|---|---|---|---|---|---|---|---|---|
| | | 1 | 2 | 3 | 4 | 5 | 6 | 7 | 8 | 9 | 10 | sum |
| Best matrix | fmgVT-1 | 10 | 6 | 6 | 6 | 9 | 9 | 1 | 9 | 9 | 2 | 67 |
| | fmgVT-2 | 5 | 10 | 7 | 4 | 2 | 2 | 5 | 2 | 6 | 5 | 48 |
| | fmgLG-3 | 9 | 8 | 10 | 5 | 4 | 7 | 7 | 5 | 7 | 4 | 66 |
| | fitVT-4 | 3 | 9 | 9 | 8 | 8 | 8 | 6 | 1 | 5 | 7 | 64 |
| | fmgLG-5 | 6 | 7 | 8 | 7 | 10 | 5 | 4 | 7 | 4 | 6 | 64 |
| | fmgVT-6 | 2 | 4 | 2 | 9 | 7 | 10 | 3 | 6 | 2 | 3 | 48 |
| | fmgVT-7 | 4 | 1 | 1 | 2 | 6 | 4 | 10 | 8 | 3 | 1 | 40 |
| | fmgVT-8 | 8 | 5 | 4 | 3 | 5 | 3 | 8 | 10 | 8 | 9 | 63 |
| | fmgLG-9 | 7 | 3 | 5 | 10 | 1 | 6 | 9 | 3 | 10 | 8 | 62 |
| | fmgVT-10 | 1 | 2 | 3 | 1 | 3 | 1 | 2 | 4 | 1 | 10 | 28 |

Supplementary File 5. Midpoint rooted Maximum likelihood tree built with CRESS. Alignment produced by MUSCLE. Green for *Geminiviridae*, teal for *Genomoviridae*, dark blue for *Smacoviridae*, red for *Bacilladnaviridae*, orange for *Circoviridae*, light blue for *Nanoviridae*, purple for the alphasatellites (*Alphasatellitidae*). Black taxa are currently unclassified.

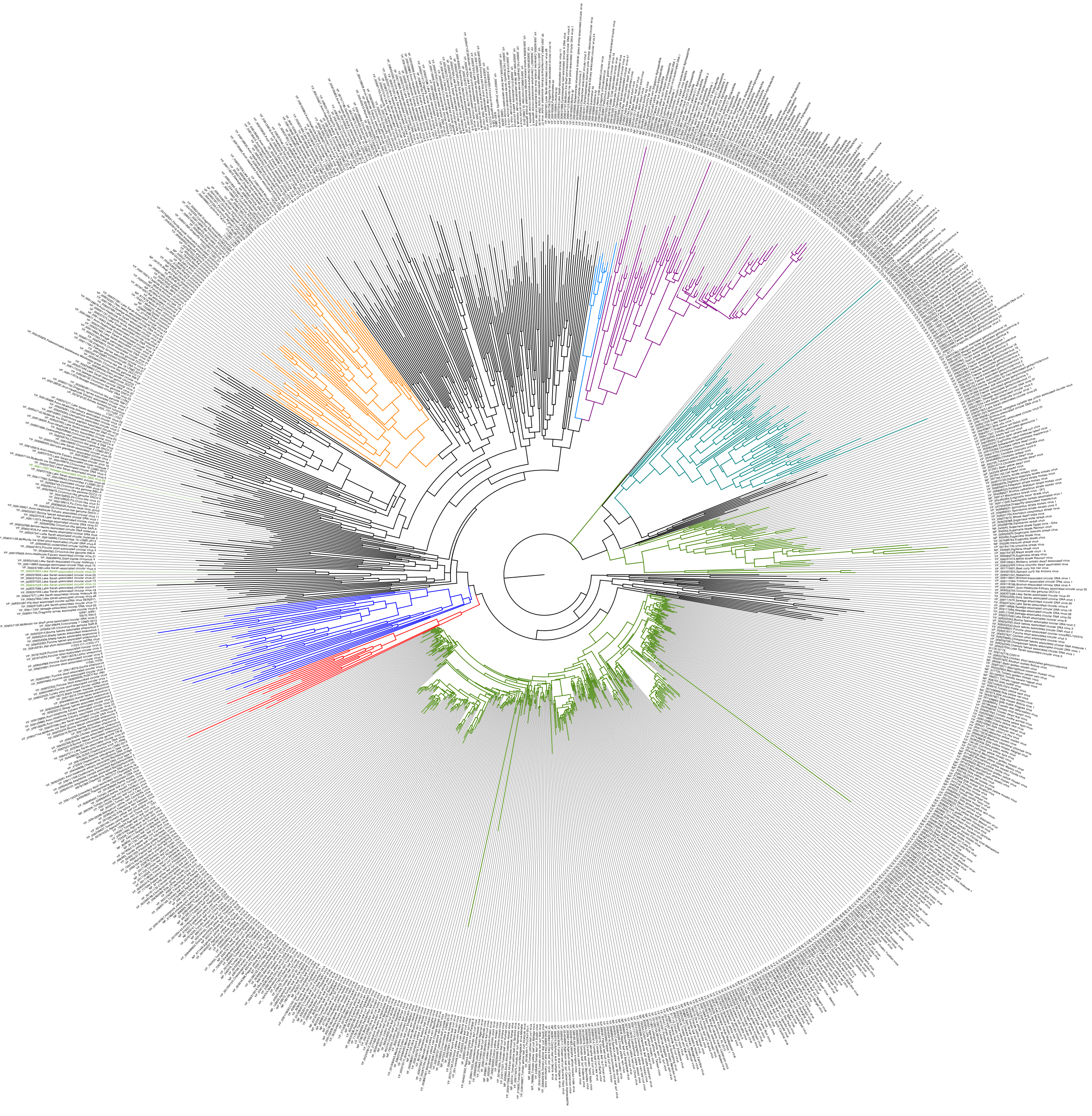

Supplementary File 7. Maximum likelihood tree built with the endonuclease portion of the full CRESS Rep MUSCLE alignment. Green for *Geminiviridae*, teal for *Genomoviridae*. Black taxa are all other taxa, both currently classified and unclassified.

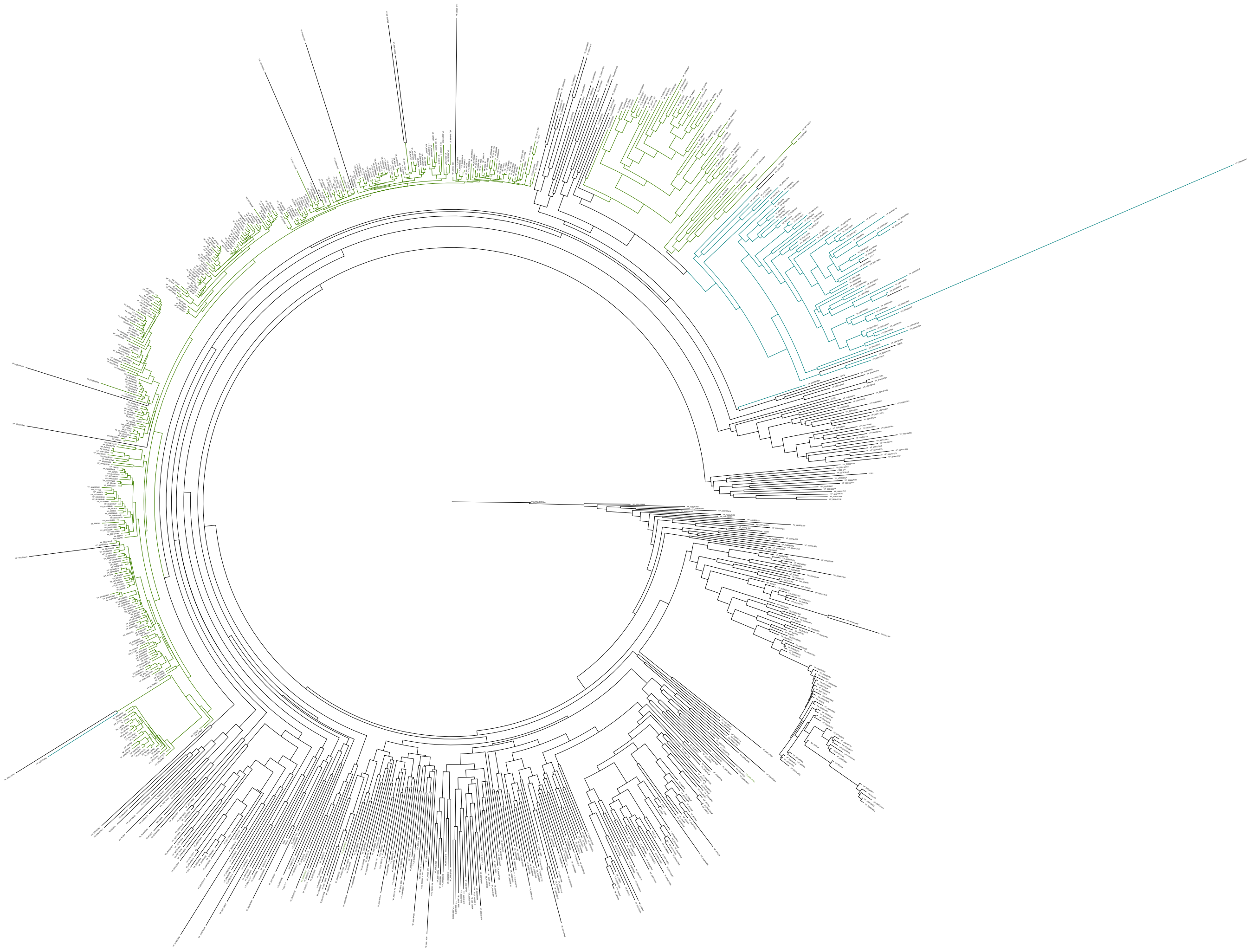

Supplementary File 8. Maximum likelihood tree built with the helicase portion of the full CRESS Rep MUSCLE alignment. Green for *Geminiviridae*, teal for *Genomoviridae*. Black taxa are all other taxa, both currently classified and unclassified.

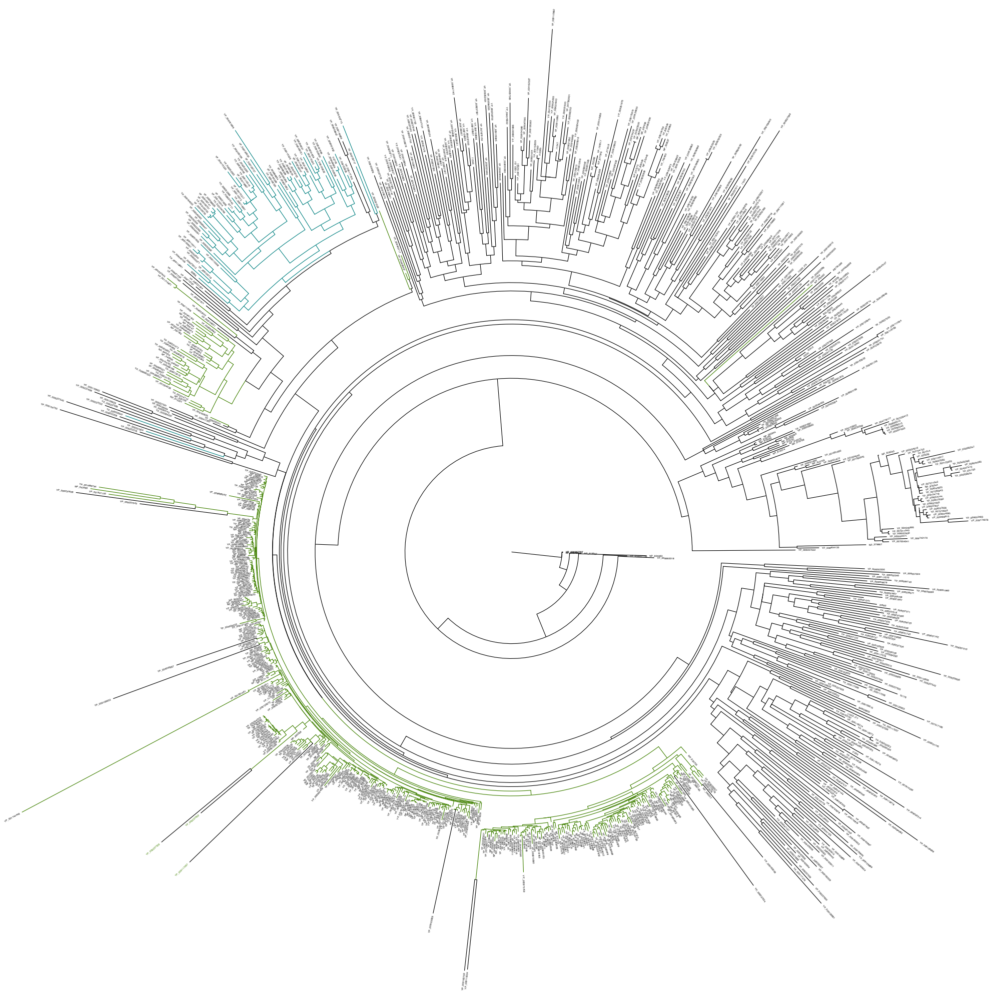

Supplementary File 9. Maximum likelihood tree with equal numbers of Begomovirus and Mastrevirus Reps built with CRESS. Alignment produced by MUSCLE. Green for *Geminiviridae*, teal for *Genomoviridae*, dark blue for *Smacoviridae*, red for *Bacilladnaviridae*, orange for *Circoviridae*, light blue for *Nanoviridae*, purple for the alphasatellites (*Alphasatellitidae*). Black taxa are currently unclassified.

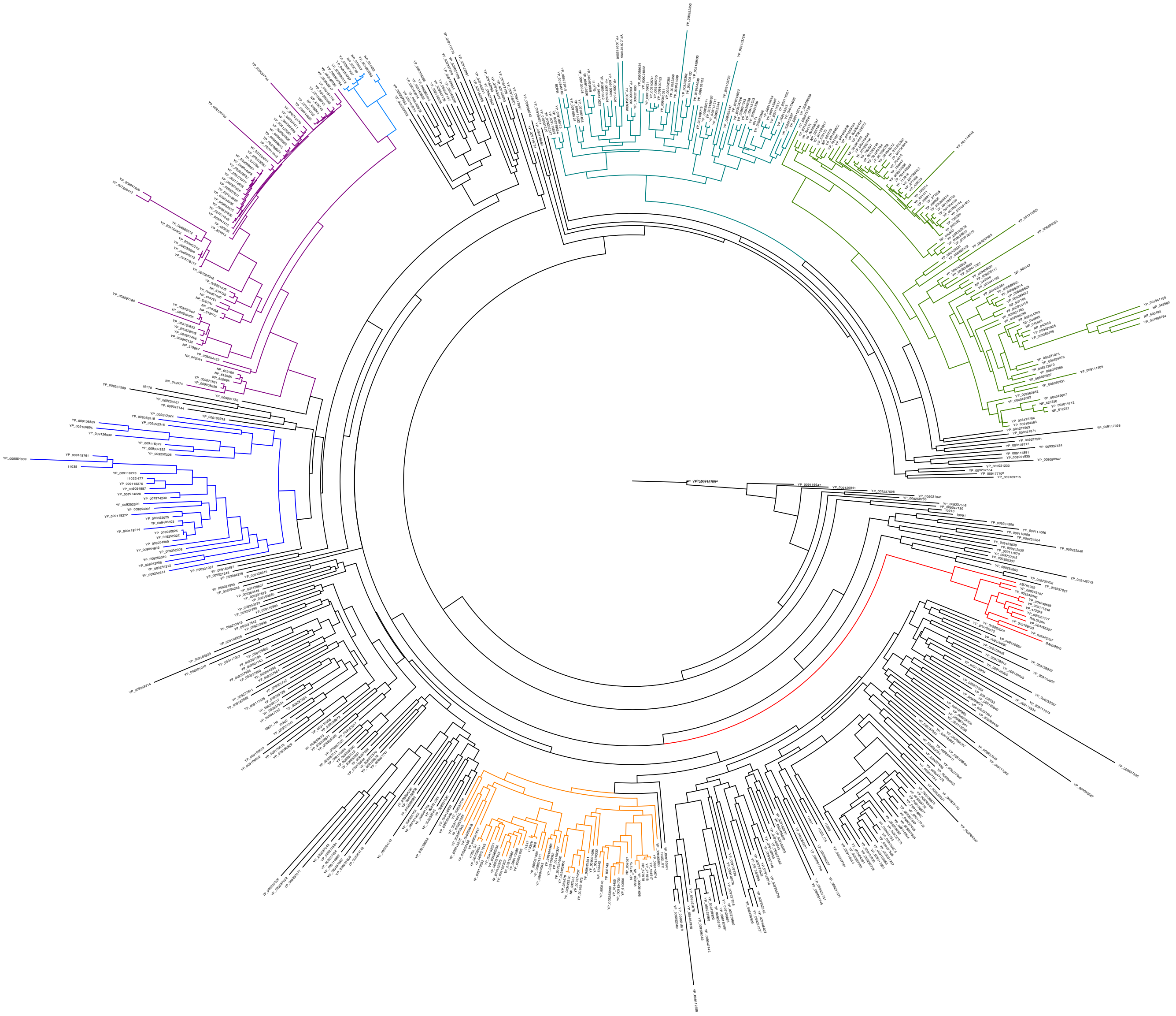

Supplementary File 10. Unrooted maximum likelihood tree built with 123 recombinant sequences removed from the 926 CRESS Rep dataset. Green for *Geminiviridae*, teal for *Genomoviridae*. Black taxa are all other taxa, both currently classified and unclassified.

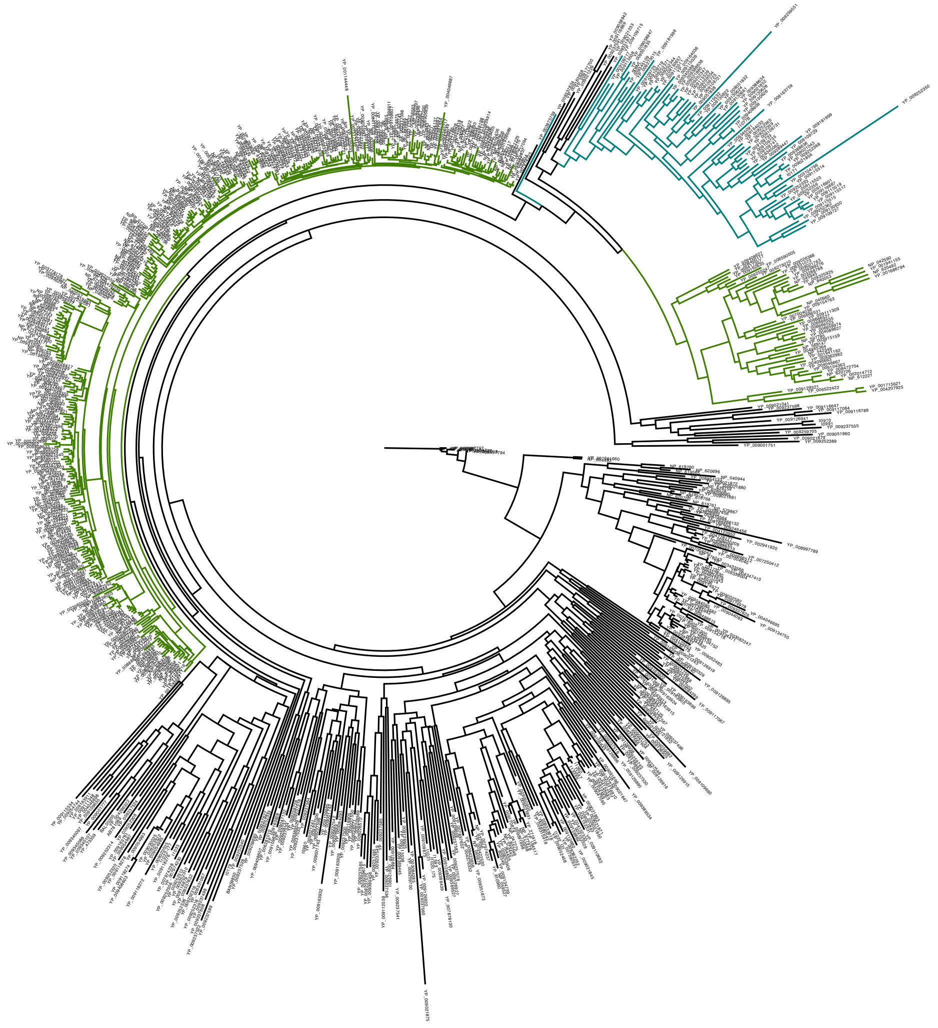

Supplementary File 11. Maximum likelihood tree built with the full 926 CRESS Rep dataset. Alignment produced by MAFFT. Green for *Geminiviridae*, teal for *Genomoviridae*, dark blue for *Smacoviridae*, red for *Bacilladnaviridae*, orange for *Circoviridae*, light blue for *Nanoviridae*, purple for the alphasatellites (*Alphasatellitidae*). Black taxa are currently unclassified.

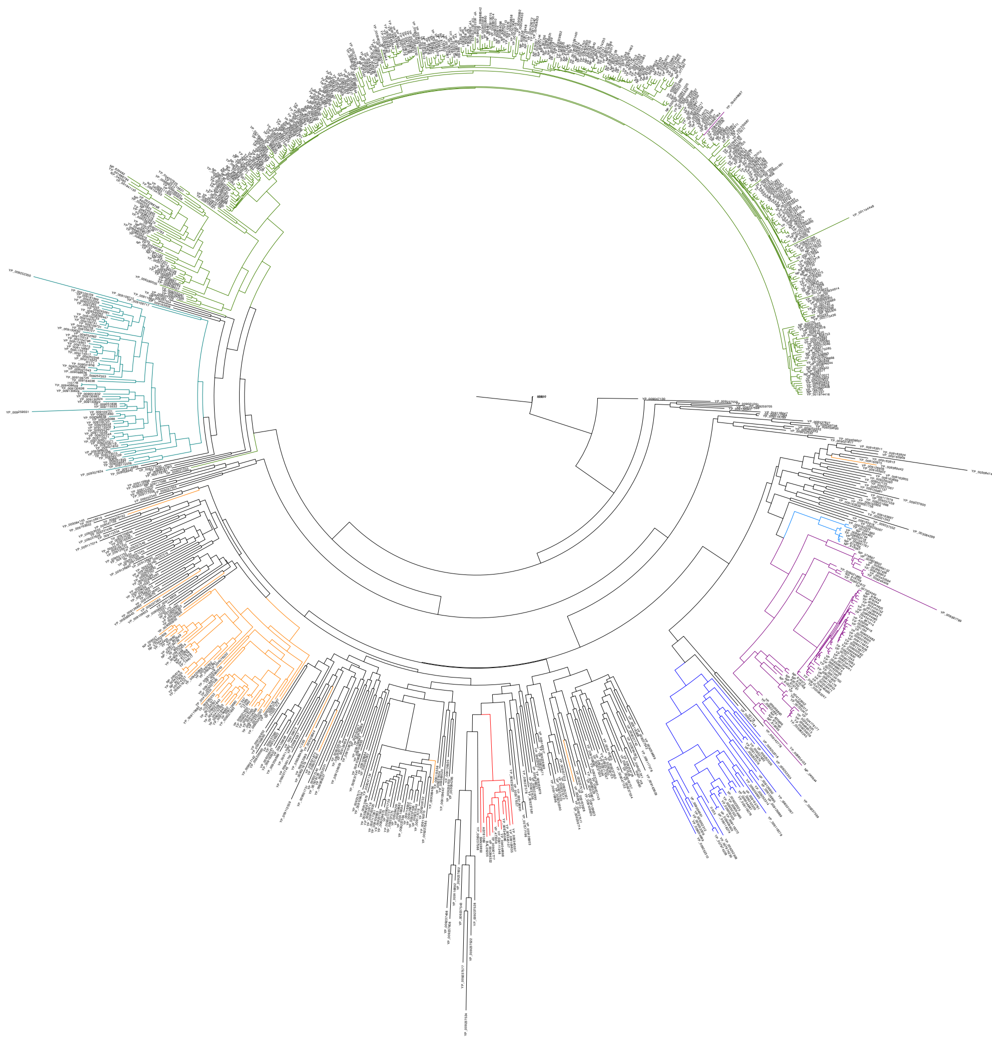

Supplementary File 12. Maximum likelihood tree with Geminivirus and Genomovirus Reps built with CRESS. Alignment product by MAFFT. Green for *Geminiviridae*, teal for *Genomoviridae*.

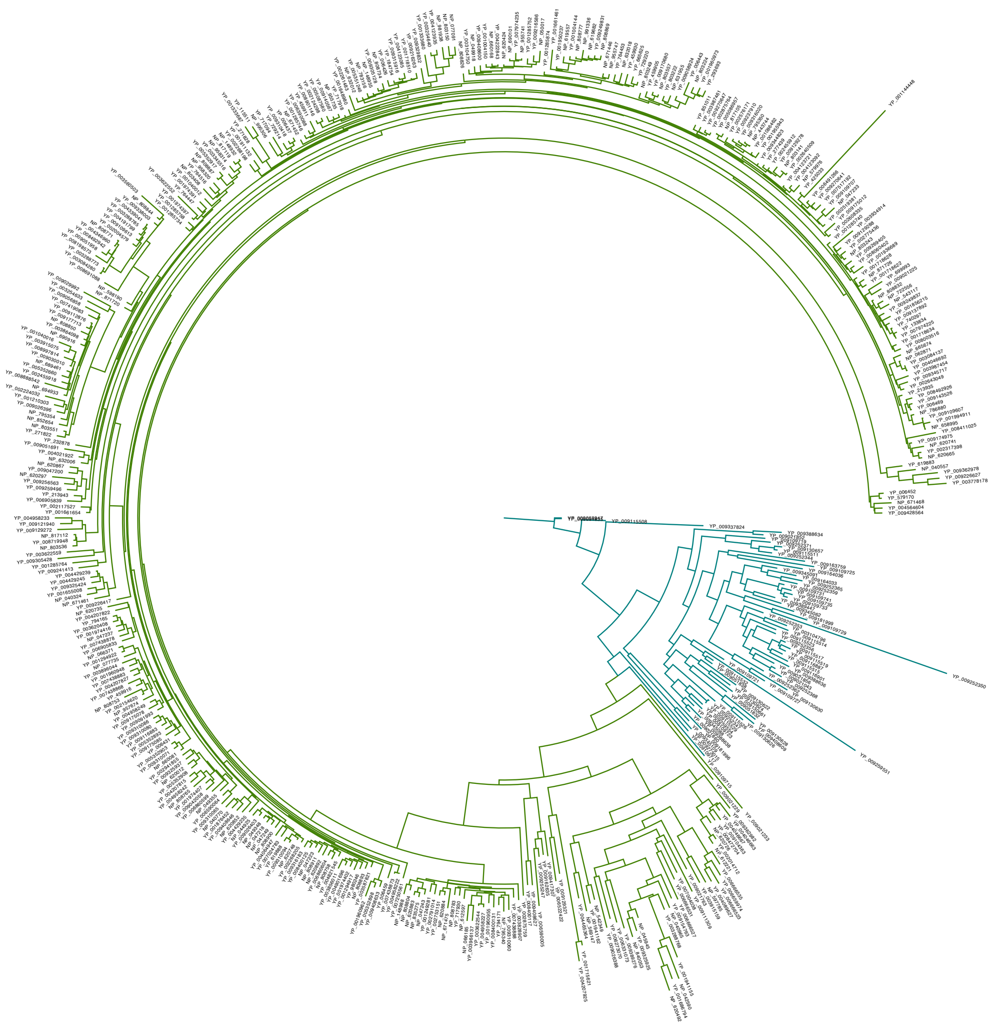

Supplementary File 13. Maximum likelihood trees with Geminivirus and Genomovirus Reps built with CRESS. Alignment produced by MUSCLE. Green for *Geminiviridae*, teal for *Genomoviridae*.

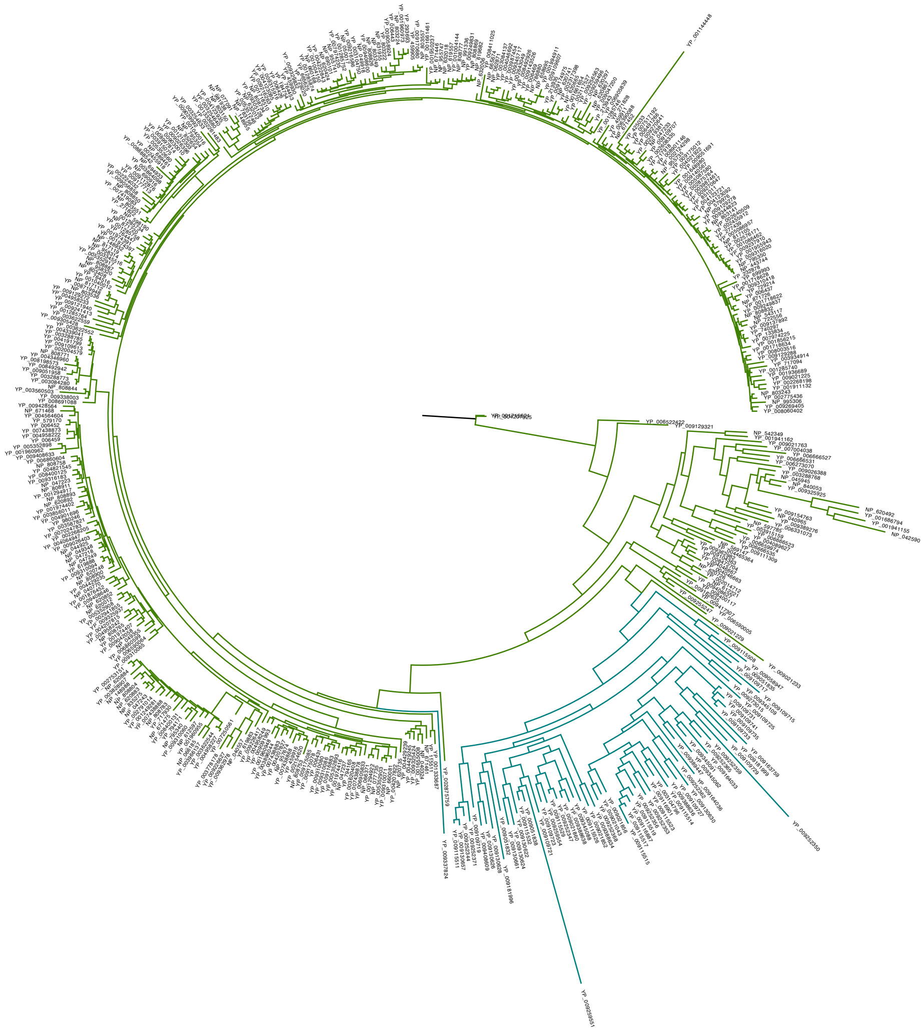

Supplementary File 14. 926 CRESS Rep sequence accession numbers. Green and blue highlighted are sequences used to build Geminivirus and Genomovirus Rep trees. Blue highlighted are removed begomoviruses in Supplement 13. Grey highlighted are collaborator provided sequences.

| | | | | | | | | | |
|---|---|---|---|---|---|---|---|---|---|
| YP_009252314 | YP_009142778 | YP_009109636 | YP_009333618 | I0831_H8 | YP_001004144 | YP_009130622 | YP_009170680 | YP_002224032 | NP_579976 |
| YP_009237555 | YP_009237588 | YP_009109649 | YP_009110682 | YP_009058942 | YP_729220 | YP_009428564 | YP_009143526 | YP_001718634 | NP_543117 |
| YP_009047130 | YP_009115534 | YP_009259720 | YP_004152327 | YP_009116889 | YP_764516 | YP_009408646 | YP_009140567 | YP_001718628 | NP_443744 |
| YP_009237524 | YP_009252340 | YP_009126916 | YP_008130363 | YP_009163938 | YP_729214 | YP_002261483 | YP_009137892 | YP_001718622 | NP_148988 |
| YP_009109686 | YP_009051960 | YP_009237546 | YP_009104366 | YP_009116891 | YP_764447 | YP_006459 | YP_009129288 | YP_006905833 | NP_148955 |
| YP_009237602 | YP_009021879 | YP_009389439 | YP_009047065 | YP_009237554 | YP_459911 | YP_009325424 | YP_009129278 | YP_003915075 | NP_148932 |
| YP_009237608 | YP_009252389 | YP_009126879 | YP_009021871 | YP_009177700 | YP_001661654 | YP_001960973 | YP_008198573 | YP_003587821 | NP_066371 |
| YP_009237498 | YP_009112559 | YP_009237585 | YP_009021850 | NP_620696 | YP_425033 | YP_009408633 | YP_003622544 | YP_003560503 | NP_062871 |
| YP_009116892 | YP_009237504 | YP_009116913 | YP_009021843 | NP_619760 | YP_001285758 | YP_009408600 | NP_047218 | YP_003345516 | NP_049918 |
| YP_009117082 | BAN59850 | YP_009259718 | YP_007392931 | NP_619565 | YP_001285740 | YP_009047200 | YP_005352908 | YP_002124298 | NP_047249 |
| YP_009117074 | AB781089 | YP_009163920 | YP_004152331 | YP_009021880 | YP_001285734 | YP_009315916 | YP_005352903 | YP_001950237 | NP_047237 |
| YP_009021243 | YP_009345097 | YP_009237575 | YP_009362252 | NP_619759 | YP_001285746 | YP_009345717 | YP_005086957 | YP_005352660 | NP_047233 |
| YP_009163897 | YP_004046698 | YP_009259682 | YP_004152333 | YP_009021872 | YP_740297 | YP_009216586 | YP_004564604 | YP_005351246 | NP_047223 |
| YP_009163932 | YP_009111348 | YP_009001737 | YP_004152329 | NP_619572 | YP_001086462 | YP_009344823 | YP_004429245 | YP_004346960 | NP_044925 |
| YP_009126901 | YP_473359 | YP_003084291 | YP_009021893 | NP_619768 | YP_277439 | YP_009338003 | YP_004429235 | YP_004339041 | NP_040770 |
| YP_009117057 | BAL05205 | YP_009109683 | YP_009110680 | NP_620700 | YP_851011 | YP_009241413 | YP_003084137 | YP_004222843 | YP_003987454 |
| YP_009237511 | YP_009001777 | YP_009117061 | YP_009021845 | NP_619761 | YP_001648980 | YP_001936689 | YP_002791014 | YP_004191799 | YP_006491266 |
| YP_009337827 | YP_004286322 | YP_009001739 | YP_009158862 | YP_009252330 | YP_717918 | YP_764453 | YP_002775436 | YP_004123092 | |
| YP_009259708 | YP_009109635 | YP_009259675 | YP_009109670 | YP_009117070 | YP_009362978 | YP_009329832 | YP_002753151 | YP_004123721 | |
| YP_009259695 | YP_009345107 | YP_009237565 | YP_009237592 | YP_009117066 | YP_009226627 | YP_009325937 | YP_002643049 | YP_004123085 | |
| YP_009237578 | YP_009345086 | YP_009237594 | YP_009109663 | YP_009237509 | YP_003778178 | YP_009316020 | YP_002608335 | YP_004046692 | |
| YP_003084285 | YP_009252327 | YP_009116894 | YP_009109643 | YP_009116898 | YP_009111309 | YP_009316183 | YP_002576171 | YP_004021922 | |
| YP_008828162 | YP_009252335 | YP_009389456 | YP_009109668 | YP_009351871 | YP_006666535 | YP_009310418 | YP_002455912 | YP_003987461 | |
| YP_009126927 | YP_009252333 | YP_009259732 | YP_009163927 | YP_003084297 | YP_006659974 | YP_009310094 | YP_002317398 | YP_003934914 | |
| YP_009109644 | YP_009237599 | YP_009126877 | YP_009117058 | YP_009116647 | YP_006666523 | YP_009310080 | YP_001974416 | YP_002268198 | |
| YP_009001747 | YP_009126930 | YP_009116902 | YP_009237577 | YP_009117064 | YP_004089627 | YP_009310065 | YP_001974402 | YP_001911132 | |
| YP_003084290 | YP_009126895 | YP_009117076 | YP_009237543 | YP_009116789 | NP_597785 | YP_009310086 | YP_001955943 | YP_003622559 | |
| YP_009109630 | YP_009126889 | YP_009237549 | YP_009126903 | YP_009126941 | YP_003915159 | YP_009310060 | YP_001876452 | YP_003288773 | |
| YP_009237570 | YP_009252324 | YP_009237495 | YP_007517186 | YP_009021041 | NP_569147 | YP_009310071 | YP_001856215 | YP_003254633 | |
| YP_009237538 | YP_009252310 | YP_009109640 | YP_009126919 | YP_009259705 | YP_004465364 | YP_009305128 | YP_980246 | YP_002875764 | |
| YP_009259692 | YP_009054989 | YP_007878130 | YP_009389445 | YP_009237598 | YP_009021763 | YP_009270641 | YP_699993 | YP_002640509 | |
| YP_009121932 | YP_009252312 | YP_009252331 | YP_009047144 | YP_009116781 | NP_542349 | YP_009270647 | YP_665625 | YP_002455918 | |
| YP_009021245 | YP_009252306 | YP_009237496 | YP_003084143 | YP_004046687 | YP_001941162 | YP_005352898 | YP_619888 | YP_001994911 | |
| YP_009126882 | YP_009252308 | YP_009259689 | YP_009160329 | YP_009237591 | YP_009026388 | YP_009269405 | YP_619883 | YP_001974397 | |
| YP_009126938 | YP_009118278 | YP_009237541 | YP_001661660 | YP_009337838 | YP_009389276 | NP_795354 | YP_232878 | YP_002268205 | |
| YP_009259745 | YP_009054991 | YP_009389527 | NP_604483 | YP_009237528 | NP_042590 | NP_620892 | YP_006469 | YP_005352917 | |
| YP_009237530 | YP_009252320 | YP_009047134 | YP_008997794 | YP_009237522 | YP_006331073 | YP_004821545 | YP_006452 | YP_005352893 | |

| | | | | | | | | |
|---|---|---|---|---|---|---|---|---|
| YP_009126890 | YP_009054987 | YP_009047142 | YP_008992018 | YP_009237563 | YP_006273070 | YP_009256563 | NP_981938 | NP_047243 |
| YP_009126896 | YP_009118276 | YP_009163907 | YP_008997797 | YP_001144448 | NP_620492 | YP_009249831 | NP_871726 | YP_459930 |
| YP_009237516 | I1022-I77 | YP_009126898 | YP_003104737 | YP_001285874 | YP_001686794 | YP_009249837 | NP_871720 | YP_001718510 |
| YP_009163905 | YP_007974230 | YP_009226561 | NP_619769 | YP_001249281 | YP_001941155 | NP_620884 | NP_852654 | YP_271822 |
| YP_009259714 | YP_007974228 | YP_009226565 | NP_619567 | YP_794171 | YP_009154763 | NP_050017 | NP_808900 | YP_213935 |
| YP_003084140 | YP_009163761 | YP_009126922 | YP_008997789 | YP_003966137 | NP_040965 | YP_002117527 | NP_808893 | YP_213943 |
| YP_009389529 | YP_009054993 | YP_009237560 | NP_579867 | NP_066185 | YP_003288768 | NP_612597 | NP_808734 | YP_184754 |
| YP_009109615 | I1035 | YP_009109660 | YP_009246456 | YP_009305428 | NP_840053 | NP_808844 | NP_803141 | YP_133834 |
| YP_009109623 | YP_009118272 | YP_009237502 | YP_003433564 | NP_040557 | YP_009325925 | NP_671446 | NP_795340 | YP_115511 |
| YP_009109620 | YP_009408603 | YP_007353980 | YP_003966132 | YP_009129272 | NP_045945 | NP_049348 | NP_722556 | YP_006426 |
| YP_003084299 | YP_009118274 | YP_009259711 | YP_003987456 | NP_620735 | YP_009362982 | YP_009237910 | NP_689461 | YP_006437 |
| YP_009021890 | YP_009022025 | YP_009126884 | YP_008169853 | YP_001294922 | YP_004046663 | YP_009226417 | NP_671461 | YP_006431 |
| YP_009237518 | YP_009054985 | YP_003084282 | YP_003828902 | YP_794165 | YP_009104363 | YP_009175085 | NP_660168 | NP_995306 |
| YP_006281010 | YP_009252322 | YP_009109626 | NP_040944 | YP_001294917 | YP_008472704 | YP_009175078 | NP_660081 | NP_991336 |
| YP_009237500 | YP_009030025 | YP_009389534 | YP_007004040 | YP_579170 | YP_004046667 | YP_003254640 | NP_632006 | NP_958320 |
| YP_009259679 | YP_009252318 | YP_009126925 | YP_008854133 | YP_001285764 | NP_620726 | YP_009216263 | NP_620883 | NP_958314 |
| YP_009226569 | YP_006331067 | YP_009259700 | YP_004123952 | YP_001210303 | YP_002014712 | NP_835275 | NP_619682 | NP_955735 |
| YP_009226571 | YP_009337832 | YP_009237586 | YP_007250412 | YP_001040016 | NP_612221 | YP_009177713 | NP_619557 | NP_955741 |
| YP_009163917 | YP_009116879 | YP_009109659 | YP_006666512 | YP_001655008 | YP_006666527 | YP_009175012 | NP_077735 | NP_955747 |
| YP_009226572 | YP_009252326 | YP_009001753 | YP_004778177 | YP_271828 | YP_007004038 | YP_009174975 | NP_077091 | NP_817119 |
| YP_009021241 | YP_009252316 | YP_009389520 | YP_009230209 | YP_009259496 | YP_006666531 | YP_006443 | YP_009116883 | NP_817112 |
| YP_009252337 | YP_009237600 | YP_009237604 | YP_002941920 | YP_001661461 | YP_009408627 | YP_009121940 | YP_008691088 | NP_817105 |
| YP_009237512 | YP_009109653 | YP_009022029 | YP_003082245 | YP_001040012 | YP_008400117 | YP_009058924 | YP_009112876 | NP_808911 |
| YP_009237514 | YP_009237571 | YP_009126892 | YP_006666513 | YP_803222 | YP_006590005 | YP_009056858 | YP_009109707 | NP_808869 |
| YP_009237520 | YP_009001745 | YP_009115538 | YP_003082247 | YP_459905 | YP_009417307 | YP_009042058 | YP_009109613 | NP_808826 |
| YP_009252390 | YP_009001751 | YP_009126935 | YP_003334471 | YP_001285752 | YP_009255247 | YP_008719948 | YP_009109607 | NP_808832 |
| YP_009116896 | YP_009117079 | YP_009126915 | YP_004046695 | YP_009021233 | YP_009162635 | YP_008400131 | YP_009091993 | NP_808850 |
| YP_009163929 | YP_009226567 | YP_009126905 | YP_004123953 | YP_001004150 | YP_009130624 | YP_008400125 | YP_009051958 | NP_808804 |
| YP_009237533 | YP_009163918 | YP_009237583 | YP_006742179 | YP_009252350 | YP_009130661 | YP_007024783 | YP_009051691 | NP_808771 |
| YP_009163928 | YP_009126881 | YP_009163901 | YP_001285945 | YP_009109715 | YP_009181999 | YP_006860604 | YP_003896044 | NP_808777 |
| YP_009001743 | YP_009226563 | YP_009109675 | YP_009154718 | YP_009337824 | YP_009252362 | YP_006860599 | YP_003622552 | NP_808758 |
| YP_009237564 | YP_009237559 | YP_009116909 | YP_009344825 | YP_009115508 | YP_009130630 | YP_006590064 | YP_003288785 | NP_808765 |
| YP_009237550 | YP_009021888 | YP_009315918 | NP_878244 | YP_009058947 | YP_009109727 | YP_004958249 | YP_003104750 | NP_808753 |
| YP_009259723 | YP_009163936 | YP_009380542 | YP_001285943 | YP_009051835 | YP_009388636 | YP_004958242 | YP_003084280 | NP_808783 |
| YP_009259728 | YP_009163904 | YP_009116910 | NP_878243 | YP_009259551 | YP_003104796 | YP_004958233 | YP_002154620 | NP_803551 |
| YP_003084293 | YP_009001756 | YP_009021847 | YP_851014 | YP_009181996 | YP_009115514 | YP_004958227 | YP_002004579 | NP_803557 |
| YP_009237506 | YP_009047125 | YP_009237526 | YP_009073582 | YP_009345109 | YP_009115523 | YP_004958222 | YP_009030010 | NP_803536 |
| YP_009237534 | YP_009047137 | YP_004376332 | YP_003104752 | YP_009273015 | YP_009021856 | YP_004901696 | YP_008411025 | NP_803408 |
| YP_009259737 | YP_009237567 | YP_164517 | YP_009254744 | YP_009163759 | YP_009021043 | YP_004429239 | YP_009029992 | NP_803243 |
| YP_009047132 | YP_009163899 | YP_271918 | YP_007004041 | YP_009388634 | YP_009252368 | YP_004207815 | YP_009026403 | NP_803249 |

| | | | | | | | | |
|---|---|---|---|---|---|---|---|---|
| YP_009117078 | YP_009163922 | NP_877978 | YP_009001905 | YP_009021852 | YP_009252353 | YP_004207827 | YP_009026396 | NP_803224 |
| YP_009116905 | YP_009126936 | YP_009091698 | YP_009044085 | YP_009164036 | YP_009252356 | YP_004207822 | YP_008997814 | NP_803150 |
| YP_009117067 | YP_009163924 | NP_150368 | YP_009052483 | YP_009109729 | YP_009116901 | YP_004123935 | YP_006390088 | NP_795350 |
| YP_009237596 | YP_009389442 | NP_047275 | YP_009134750 | YP_009109725 | YP_009115515 | YP_004064947 | YP_009021225 | NP_786880 |
| YP_009259699 | YP_009163909 | YP_803546 | YP_425036 | YP_009130628 | YP_009115519 | YP_003856011 | YP_008492942 | NP_783156 |
| YP_009237542 | YP_009163913 | NP_059527 | YP_004347413 | YP_009130626 | YP_009116887 | YP_003828907 | YP_008901146 | NP_694933 |
| YP_009001742 | YP_009163911 | NP_573442 | YP_009337826 | YP_009408609 | YP_009115517 | YP_003620408 | YP_003864098 | NP_690916 |
| YP_009109685 | YP_009163915 | YP_610960 | YP_009174978 | YP_009109719 | YP_009164033 | YP_002941855 | YP_001974391 | NP_690101 |
| YP_009116906 | YP_009109676 | YP_009134739 | YP_009174977 | YP_009252371 | YP_009345091 | YP_002519381 | YP_008888542 | NP_671468 |
| I0178 | YP_009021875 | YP_803549 | YP_009002582 | YP_009252344 | YP_009252365 | YP_001960948 | YP_008492926 | NP_671475 |
| I1020_I75 | YP_009001750 | YP_764455 | YP_009044083 | YP_009130657 | YP_009252359 | YP_001960955 | YP_007974235 | NP_671452 |
| I1021 | YP_003084287 | YP_009253909 | YP_009337831 | YP_009115511 | YP_009109731 | YP_001960962 | YP_007974225 | NP_665674 |
| I1393 | YP_009126929 | YP_009091696 | YP_007518505 | YP_009115526 | YP_009109741 | YP_001974407 | YP_005087580 | NP_658995 |
| I1022_I72 | YP_009259555 | YP_009170674 | YP_003934916 | YP_009345088 | YP_009109735 | YP_001333680 | YP_008060402 | NP_632018 |
| I1022_G1 | YP_009047139 | YP_009351873 | YP_031730 | YP_009388638 | YP_009109733 | YP_001333687 | YP_008003516 | NP_632012 |
| I1360 | YP_009408607 | YP_007974237 | YP_007011043 | YP_009021860 | YP_009389447 | YP_717930 | YP_007517192 | NP_631955 |
| I1338b | YP_009021877 | YP_003422530 | YP_009337830 | YP_009252347 | YP_009345062 | YP_717094 | YP_007438888 | NP_620852 |
| I1347 | YP_009237581 | NP_065678 | YP_007011042 | YP_009115529 | YP_001715621 | YP_459916 | YP_007438883 | NP_620741 |
| SDBVL | YP_009237552 | NP_937956 | YP_006488615 | YP_009259554 | YP_004207925 | YP_293693 | YP_007438878 | NP_620748 |
| I1077B | YP_009126932 | YP_009021891 | YP_003433566 | YP_009109723 | YP_009021229 | NP_957674 | YP_007438873 | NP_620665 |
| I0171 | YP_009237536 | YP_009423856 | YP_009338004 | YP_009051832 | YP_009109717 | NP_808887 | YP_007438868 | NP_620424 |
| I0910 | YP_009237606 | YP_007697652 | NP_619574 | YP_009109721 | YP_009129321 | NP_620867 | YP_007419083 | NP_620297 |
| I0991 | YP_009252342 | YP_009000900 | YP_009058890 | YP_009115532 | YP_006522422 | NP_049355 | YP_006905839 | NP_620012 |
| I0960 | YP_009237540 | YP_009052458 | YP_009021881 | YP_009051838 | YP_002875759 | NP_040324 | YP_007250561 | NP_598190 |

Additional manual edits: YP_009408627 and YP_009408626 were concatenated by removing amino acids 207-261 from YP_009408627, amino acids 1-21 from YP_009408626. YP_008400117 and YP_008400116 were concatenated by removing amino acids 205-264 from YP_009408627, amino acids 1-16 from YP_009408626. NP_040965 was NP_040964 and NP_040965 concatenated with the removal of amino acids 214-266 from NP_040965. NP_045945 was NP_045945 and NP_045944 concatenated with the removal of amino acids 224-276 from NP_045945. NP_597785 was NP_597785 and NP_899201 concatenated with the removal of amino acids 272-336 from NP_587785. NP_840053 was NP_840053 and NP_840052 concatenated with the removal of amino acids 219-312 from NP_840053. YP_006522422 was YP_006522422 and YP_006522421 concatenated with the removal of amino acids 205-269 from YP_006522422. Other concatenated sequences include NP_569147+46, YP_001716521+20, YP_004207925+24, without removal of amino acids from sequence.

Supplementary File 16. Genome accession numbers from which capsid protein sequences were extracted.

| Genomovirus | | Circovirus | Smacovirus | Nanovirus | Bacilladnavirus | Parvovirus | |
|---|---|---|---|---|---|---|---|
| KT862253 | KF371630 | KM382269 | KM573772 | Y003104738 | AB597949 | NC_004285 | NC_035186 |
| KR912221 | KF371631 | KJ641740 | KT862221 | N619570 | AB844272 | NC_015115 | NC_016647 |
| KT732792 | LK931484 | KT732785 | KT862225 | Y008997806 | AB193315 | NC_011317 | NC_023673 |
| KT732793 | KP974693 | KT732786 | KT862219 | Y008997802 | AB553581 | NC_012636 | NC_016031 |
| KF371640 | KJ547634 | KM017740 | KT862223 | Y008992019 | AB781089 | NC_004290 | NC_016032 |
| KT862251 | KJ938717 | JX185424 | KM573769 | N619767 | KY405008 | NC_022748 | NC_012042 |
| KF371643 | KP263543 | KT732787 | KT862224 | N620699 | KF133809 | NC_004289 | NC_012729 |
| KF371641 | LK931483 | KC771281 | KT862218 | Y009508036 | KY405006 | NC_015718 | NC_012564 |
| KF371642 | KP263545 | KF031466 | KT862222 | N604477 | KY405007 | NC_006555 | NC_007455 |
| GQ365709 | KP263546 | GQ404857 | JN634851 | Y001661657 | | NC_018450 | NC_014358 |
| KF268025 | KP987887 | HQ738634 | KM598409 | Y009508220 | | NC_004287 | NC_031695 |
| KF268026 | KT862250 | KJ641712 | KT862228 | | | NC_018399 | NC_029133 |
| KF268027 | KP133076 | JF938079 | KM573774 | | | NC_004288 | NC_028973 |
| KF268028 | KP133077 | GQ404846 | KP233194 | | | NC_023842 | NC_024453 |
| KM598382 | KP133078 | GQ404845 | KT600068 | | | NC_005040 | NC_029300 |
| KM598383 | KP133079 | KJ831064 | GQ351272 | | | NC_012685 | NC_030873 |
| KM598384 | KP133080 | HQ738643 | GQ351275 | | | NC_001899 | NC_022800 |
| KF371639 | KP133075 | HQ738637 | KJ577810 | | | NC_019492 | NC_017823 |
| KT253577 | KP263547 | GQ404849 | KJ577817 | | | NC_004286 | NC_020499 |
| KT253578 | KJ641737 | JF938081 | KF880727 | | | NC_005341 | NC_004442 |
| KT253579 | KP263544 | KC512919 | KJ577819 | | | NC_000936 | NC_025825 |
| KF371637 | HQ335086 | HQ738636 | KM573770 | | | NC_030296 | NC_031751 |
| JX185429 | JX185430 | JX185426 | KC545226 | | | NC_026943 | NC_034445 |
| KM598385 | KT862249 | JF938082 | KC545227 | | | NC_005041 | NC_001662 |
| KM598386 | KJ641726 | HM228874 | KP233191 | | | NC_031450 | NC_001718 |
| KM598387 | KT732790 | JX569794 | KP233178 | | | NC_014357 | NC_001510 |
| KM598388 | KT732791 | GQ404844 | KP233180 | | | NC_007218 | NC_001539 |
| KT732801 | KJ413144 | KC512920 | KY086301 | | | NC_011545 | NC_029797 |
| KT732802 | KJ547635 | KJ641715 | KP233174 | | | NC_032097 | NC_024888 |
| KF371635 | KT862254 | KJ641720 | KY086300 | | | NC_004284 | NC_030837 |
| JQ412056 | KT862238 | KM382270 | KP860906 | | | NC_002190 | NC_028650 |
| JQ412057 | KT862239 | GQ404854 | KT862229 | | | NC_022564 | NC_026815 |
| KT732794 | JN704610 | JX185422 | KP233190 | | | NC_022089 | |
| KF371633 | KJ547642 | KJ641717 | JX274036 | | | NC_037053 | |
| KT862244 | KT732813 | KF726984 | KJ577811 | | | NC_000883 | |
| KT862243 | KJ547644 | KJ641728 | KM573771 | | | NC_004295 | |
| KT862246 | KP974694 | GQ404855 | KM573775 | | | NC_026251 | |
| KT363839 | KJ547645 | JX185419 | KJ577816 | | | NC_016752 | |

| | | | | | | |
|---|---|---|---|---|---|---|
| KT732795 | LK931485 | KC512918 | KP233189 | | | NC_006259 |
| KT732796 | KM510192 | LC018134 | KM598410 | | | NC_031959 |
| KJ641719 | KT732814 | GQ404847 | KT862220 | | | NC_023860 |
| KT732800 | KJ547639 | KC512916 | KJ577812 | | | NC_014665 |
| KT732806 | KJ547637 | KR902499 | GQ351273 | | | NC_023020 |
| KT732804 | KJ547640 | KC241982 | | | | NC_027429 |
| KT732805 | KJ547638 | KJ641711 | | | | NC_006148 |
| JX185428 | KM821747 | KJ641727 | | | | NC_006147 |
| KT862242 | KT862245 | AF071878 | | | | NC_001701 |
| KT309029 | KT862247 | AF252610 | | | | NC_014468 |
| KT732798 | KF371634 | DQ172906 | | | | NC_006263 |
| KT732799 | KJ547643 | GQ404851 | | | | NC_004828 |
| KT732808 | KT862255 | DQ845074 | | | | NC_006261 |
| KT732812 | | AJ301633 | | | | NC_002077 |
| KT732810 | | DQ146997 | | | | NC_006260 |
| KT732811 | | DQ845075 | | | | NC_001729 |
| KT732807 | | KP793918 | | | | NC_001829 |
| KT732809 | | GU799606 | | | | NC_005889 |
| KT732797 | | DQ100076 | | | | NC_006152 |
| KF413620 | | AJ304456 | | | | NC_025891 |
| KT732803 | | EU056309 | | | | NC_035180 |
| KF371632 | | JQ814849 | | | | NC_025965 |
| KT598248 | | KT783484 | | | | NC_016744 |
| KF371638 | | KC339249 | | | | NC_007018 |
| KT862241 | | AF027217 | | | | NC_022104 |
| KT862240 | | AF071879 | | | | NC_031670 |
| KT862252 | | JX863737 | | | | NC_028136 |
| KU343137 | | KJ020099 | | | | NC_024452 |
| KJ547641 | | JQ011377 | | | | NC_024454 |
| KJ547636 | | GQ404856 | | | | NC_030402 |
| KF371636 | | KJ641723 | | | | NC_001540 |
| KT862248 | | KJ641724 | | | | NC_035185 |
